# Genomic and phenotypic evolution of *Escherichia coli* in a novel citrate-only resource environment

**DOI:** 10.1101/2020.01.22.915975

**Authors:** Zachary D. Blount, Rohan Maddamsetti, Nkrumah A. Grant, Sumaya T. Ahmed, Tanush Jagdish, Brooke A. Sommerfeld, Alice Tillman, Jeremy Moore, Joan L. Slonczewski, Jeffrey E. Barrick, Richard E. Lenski

## Abstract

Evolutionary innovations allow populations to colonize new, previously inaccessible ecological niches. We previously reported that aerobic growth on citrate (Cit^+^) evolved in a population of *Escherichia coli* during adaptation to a minimal glucose medium containing citrate (DM25). Cit^+^ can grow in citrate-only medium (DM0), which is a novel environment for *E. coli*. To study adaptation to this new niche, we evolved one set of Cit^+^ populations for 2,500 generations in DM0 and a control set in DM25. We identified numerous parallel mutations, many mediated by transposable elements. Several lineages evolved multi-copy amplifications containing the *maeA* gene, constituting up to ∼15% of the genome. We also found substantial cell death in ancestral and evolved clones. Our results demonstrate the importance of copy-number variation and transposable elements in the refinement of the Cit^+^ trait. However, the observed mortality suggests a persistent evolutionary mismatch between *E. coli* physiology and a citrate-only environment.

## INTRODUCTION

Evolutionary novelties are qualitatively new traits that allow populations to invade previously inaccessible ecological niches (Simpson 1953, Mayr 1960). Novel traits are thus important drivers of speciation and adaptive radiations that promote biodiversity and ecological complexity. Indeed, many of the major transitions in evolution have been mediated by novel traits such as photosynthesis, multicellularity, endoskeletons, sociality, and cognition (Maynard Smith and Szathmary 1997, Lundgren et al. 2015, Erwin 2017, 2019).

We previously proposed a model in which novel traits evolve via three distinct phases (Blount et al. 2012). In the potentiation phase, mutations accumulate in a lineage that make it possible to evolve the trait. In the actualization phase, a specific mutation produces the trait. Newly evolved traits are typically weak and ineffective. However, if the new trait confers even a slight advantage, it may spread through a population and, in the refinement phase, be improved by natural selection acting on subsequent mutations.

While potentiation and actualization enable the emergence of a novel trait, the capacity for refinement affects the trait’s long-term persistence and potential to influence subsequent evolution (Quandt et al. 2015, Erwin 2015). Prospects for refinement depend on the capacity to generate heritable phenotypic variation that can improve the trait and integrate it with other aspects of organismal performance (Kirshner and Gerhart 1998; Pigliucci 2008), a facet of evolvability that we call “refinement potential.”

Refinement potential is likely crucial for a population’s long-term success in a new niche. A novel trait might allow a lineage to discover a new niche, but it does not guarantee long-term persistence. The new conditions may expose the population to selection pressures that differ in important respects from those of its ancestral niche, resulting in a mismatch between the organism and its environment (Yeh 2004, Schluter and Conte 2009, Hu et al. 2017). Successful establishment may thus depend on ameliorating this mismatch (Chang et al. 2011, Turkarslan et al. 2011), and failure to further adapt may lead to invasion failures (Zenni and Nuñez 2013). Adaptation to a new niche therefore requires a fine balance between evolvability and robustness (Lenski et al. 2006). The benefits of refining a novel trait has to outweigh the costs (if any) of integrating that trait into organismal physiology.

Adaptation to novel niches has been widely studied in the context of invasive species that colonize and adapt to unfamiliar environments (Davis 2009, MacDougall et al. 2009, Logan et al. 2019). However, the ongoing refinement of traits that provide access to novel niches has received little attention, probably because most evolutionary novelties (and associated niche discoveries) occurred in the distant past and are therefore difficult to study. Experimental evolution allows researchers to overcome this challenge. It is possible to study evolutionary novelties that arise during experiments with microbial (Blount et al. 2008; Ratcliff et al. 2012, Barrick and Lenski 2013, Kassen 2019) and digital (Lenski et al. 2003) systems, in which evolution can be studied in real-time.

One such system is the Long-Term Evolution Experiment with *Escherichia coli* (LTEE), in which 12 bacterial populations founded from a common ancestral strain have been propagated for >70,000 generations in a glucose-limited minimal medium, DM25 (Lenski et al. 1991). DM25 also contains abundant citrate, which serves as an iron-chelating agent (Blount 2016). Many bacteria can grow aerobically on citrate, but most *E. coli* cannot due to an inability to transport citrate into the cell (Koser 1924, Hall 1982, Reynolds and Silver 1983, Pos et al. 1998).

Citrate went unexploited as a nutrient in all of the LTEE populations until a Cit^+^ variant evolved in the population designated Ara–3 after ∼31,000 generations (Blount et al. 2008). The Cit^+^ phenotype arose by a genetic duplication that activated a previously unexpressed di- and tricarboxylate transporter (Blount et al. 2012). The benefit of this duplication mutation was contingent, at least in part, on that lineage’s prior evolution of an enhanced ability to use acetate excreted into the medium as a byproduct of glucose metabolism. That enhanced ability resulted from a mutation in citrate synthase that altered carbon flow into the tricarboxylic acid cycle in a manner that was pre-adaptive for growth on citrate (Quandt et al. 2015). Concurrently, the supply of competing beneficial mutations of large effect declined later in the LTEE, allowing the Cit^+^ lineage to escape competitive exclusion (Leon et al. 2018). The Cit^+^ trait radically altered this population’s ecology and subsequent evolution (Blount et al. 2008, Blount et al. 2012, Quandt et al. 2015, Quandt et al. 2014, Turner 2015, Turner et al. 2015). Access to the large citrate pool in the medium led to a several-fold increase in population size (Blount et al. 2008). Nonetheless, a Cit^−^ lineage stably coexisted with the new Cit^+^ lineage for some 10,000 generations, before finally going extinct (Blount et al. 2008, Blount et al 2012, Turner 2015, Turner et al. 2015). Evan after 70,000 generations, none of the other 11 populations in the LTEE have evolved the ability to use the available citrate (Blount et al. 2019).

The emergence of Cit^+^ in the LTEE provides a powerful model system to study the process of evolutionary innovation. Cit^+^ variants can grow not only in DM25, which contains both glucose and citrate, but also in DM0, a citrate-only medium. How would the Cit^+^ trait be refined if these variants colonized and adapted to this newly accessible citrate-only environment? To address this question, we founded 12 new, initially clonal Cit^+^ populations and allowed them to evolve in DM0 for 2,500 generations. We also evolved a second set of 12 populations in the original DM25 medium. Here we present several analyses of these populations and their evolution. We compared the growth of the DM0-evolved clones to the growth of their ancestors in both DM0 and DM25. We sequenced the genomes of evolved clones sampled from all 24 populations, which allowed us to find parallel genetic changes that identify likely targets of selection (Tenaillon et al. 2016, Deatherage et al. 2017). Among other parallel changes, we identified numerous IS element insertions and several large gene amplifications. Our results show that genomic structural variation involving transposable elements and amplifications can provide a rich source of plasticity and potential for adaptation to novel niches. Though all populations show substantial adaptation, we found evidence of persistent maladaptation, too. Some individual clones grow more poorly than their ancestors in the medium in which they evolved. We also observed high levels of cell death in both the ancestral and evolved Cit^+^ clones, highlighting underlying metabolic challenges associated with this new function. This experimental system thus sheds light not only on how new traits are refined during adaptation to a novel niche, but also on how maladaptive phenotypes may persist for long periods in new environments.

## RESULTS

### Experimental design and phylogenetic analysis of sequenced strains

We isolated three Cit^+^ clones (CZB151, CZB152, and CZB154) from the 33,000-generation sample of the Ara–3 population, and isolated spontaneous Ara^+^ revertants for each (ZDB67, ZDB68, and ZDB69, respectively). We used each of the six clones to found two populations that evolved in the citrate-only medium (DM0) for 2,500 generations and two populations that evolved for 2,500 generations in the medium containing both glucose and citrate (DM25) as a control. Evolved clones were isolated from each of the 24 populations at the end of the experiment, and we sequenced their genomes along with those of the six founding clones. We used these data to identify mutations that had accumulated during the evolution experiment.

We also used the genomic data to verify the presumed phylogenetic relationships among the ancestral (including the Ara^+^ revertants) and evolved clones, in the context of the Cit^+^ lineage of the Ara–3 population. This analysis showed that CZB154 is one mutation off the line of descent for the post-33,000 generation Cit^+^ lineage in the Ara–3 population, as it subsequently evolved in the LTEE (Blount et al. 2012). That mutation is a 1-bp deletion (GGGGGG → GGGGG) in the promoter of the hypothetical protein-coding gene ECB_03525. CZB151 does not have that mutation, but it possesses all of the other mutations found in the CZB154 clone, as well as two additional mutations. One is a C→G transversion that causes a nonsynonymous E181K mutation in the *insD* transposase. The other is a deletion of a CGCGG repeat that restores both the reading frame and function to the pseudogene *dcuS* (Turner 2015). The restored gene encodes a histidine kinase that regulates anaerobic fumarate respiration (UniProt Consortium 2017). CZB152, by contrast, belongs to a lineage somewhat further from the eventual line of Cit^+^ descent in the Ara–3 population, and it differs from CZB151 and CZB154 by several mutations (Blount et al. 2012). Genomic analysis also showed that the Ara^+^ revertant ZDB67 differs from its parent clone, CZB151, only in the expected restoration-of-function mutation in the *araA* gene. The Ara^+^ revertants ZDB68 and ZDB69, by contrast, possess secondary mutations relative to CZB152 and CZB154, respectively, in addition to the expected mutation in *araA*. ZDB68 has a C→G transversion that introduces a nonsynonymous T33I mutation in *yfcC*, which encodes a predicted inner-membrane protein; and ZDB69 has a 1-bp deletion in *nplI,* which encodes a hypothetical protein of unknown function. All 24 evolved clones evolved in DM0 or DM25 have the mutations that are unique to their ancestors. Therefore, no cross-contamination that would compromise the independence of the evolved lines took place during the experiment.

One evolved clone from population DM0–2, ZDBp874, lacked the *citT* amplification that confers the Cit^+^ phenotype (Blount et al. 2008, Blount et al. 2012). The clone displayed a negative reaction on Christensen’s Citrate agar, confirming a Cit^−^ phenotype. We therefore sequenced the genome of a second isolate, ZDBp875, from the same population. ZDBp875 was verified as having the Cit^+^ phenotype. Comparison of the ZDBp874 and ZDBp875 genomes showed that they share a single derived mutation, an IS*150* insertion at the –35 position of the promoter of *yhiO*, which encodes the putative universal stress protein B (UspB). The two clones thus appear to belong to coexisting lineages that diverged early during the evolution in DM0. We did not discover any additional Cit^−^ variants in the DM0–2 population during a phenotypic screen of several hundred clones. Previous work has shown that the *citT* amplifications are prone to spontaneous collapse back to a single copy (Blount et al. 2012). This collapse presumably occurs by homologous recombination. In any case, it eliminates CitT expression and causes reversion to the Cit^−^ phenotype. Thus, the ZDBp874 clone appears to be a recent and fortuitously sampled “amplification collapse” mutant, rather than a representative of a stably coexisting Cit^−^ lineage in that population.

### Genome evolution is faster in the citrate-only environment than in the control environment

The populations that evolved in the citrate-only DM0 medium accumulated more mutations than those that evolved in DM25, which contains both glucose and citrate (Fig. 1A vs. Fig. 1B). The DM0-evolved genomes had an average of 19.5 mutations, whereas the DM25-evolved genomes had an average of 13.7 mutations (Mann-Whitney two-tailed test, *p* = 0.0116). The DM0 genomes had an average of 3.1 nonsynonymous SNPs in protein-coding genes, as compared to 1.1 on average for the DM25 genomes. The DM0 genomes also had more IS insertions on average than the DM25 genomes (10.3 vs. 6.6), driven largely by IS*150* insertions (8.5 vs. 4.9). The evolved genomes from the DM0 and DM25 treatments had similarly low average numbers of synonymous mutations (0.1 vs. 0.3), deletions (3.8 vs. 3.8), non-IS insertions (0.8 vs. 0.8), consecutive bp-substitutions (0.1 vs. 0.1), and SNPs outside of coding regions (1.4 vs. 1.1). We cannot tell whether the large difference in IS insertions reflects higher transposition rates, stronger selection, more cell divisions resulting from differences in death rates (Frenoy and Bonhoeffer 2018), or some combination thereof. By contrast, the nearly identical numbers of synonymous mutations and SNPs outside protein-coding genes implies that the disparity in nonsynonymous mutations between the two conditions was driven by more intense selection in the DM0 environment.

**Figure 1.**
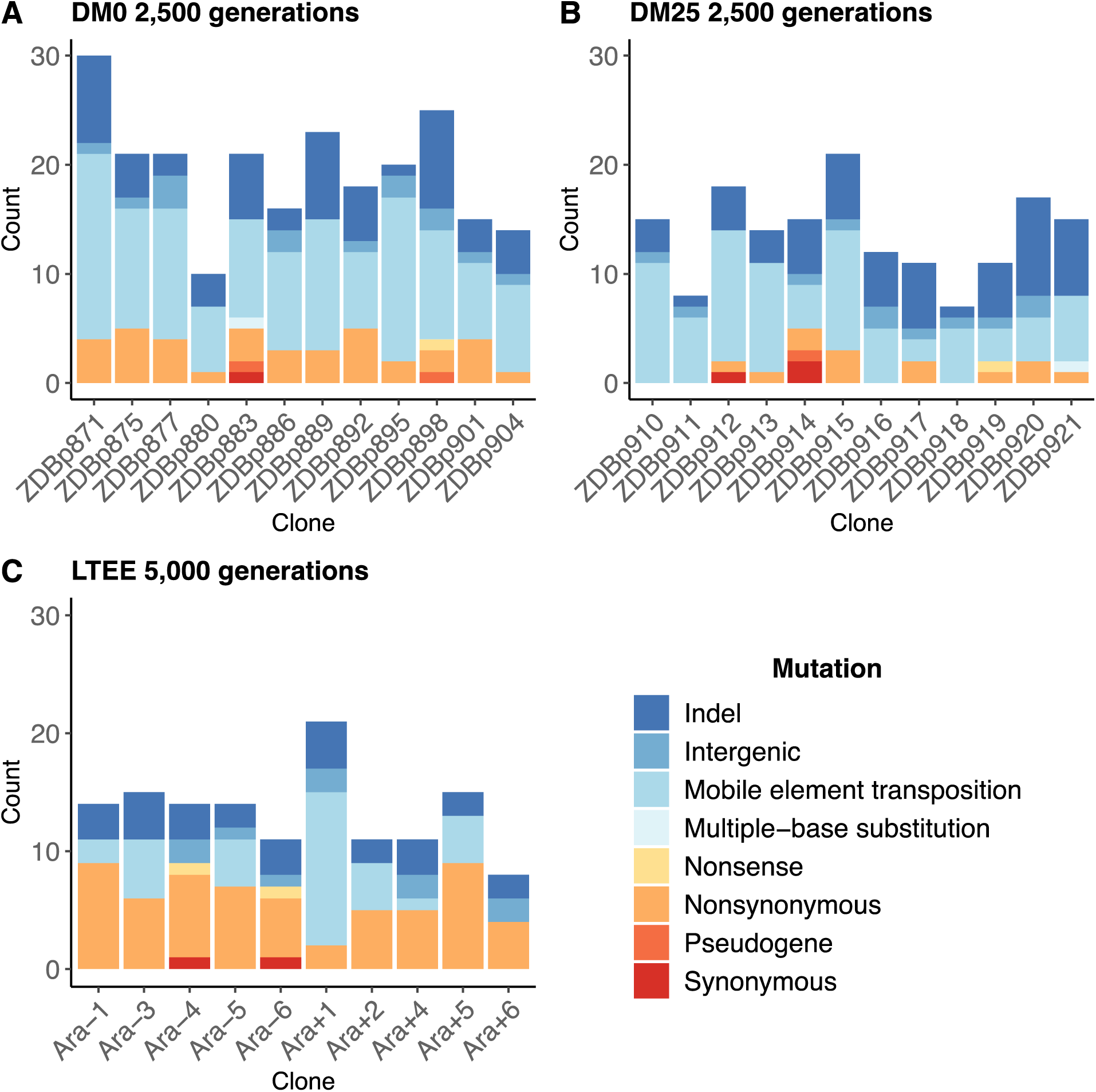
Numbers and types of mutations in evolved genomes. A) Evolved genomes from the DM0 treatment after 2,500 generations. B) Evolved genomes from the DM25 treatment after 2,500 generations. C) Evolved genomes in the 10 non-hypermutable LTEE populations after 5,000 generations. Mutations are color-coded according to the key: Indel, insertions and deletions (excluding large duplications and amplifications); intergenic, intergenic point mutations; mobile-element transpositions; multiple-base substitution, consecutive point mutations (including adjacent to and in conjunction with indels); nonsense, nonsynonymous, and synonymous point mutations in protein-coding genes; pseudogene, mutations in pseudogenes.

In both resource environments, the spectrum of mutations identified in evolved clones was dominated by structural variation, including insertions, deletions, and mobile element transpositions (Fig. 1A, B). This spectrum is very different from that observed in the LTEE populations. Figure 1C shows the number and spectrum of mutations in clones isolated from 10 LTEE populations after 5,000 generations (two other populations had evolved point-mutation hypermutability and are not shown). Despite having evolved for twice as many generations, the LTEE clones had roughly similar numbers of mutations as observed in our experiments. The mutational spectrum was dominated by nonsynonymous point mutations in all but one of the LTEE populations, Ara+1. The mutation spectrum in our study is similar to that particular population, which evolved an elevated rate of IS*150* transposition early in its history (Papadopoulos et al. 1999, Tenaillon et al. 2016). It is also similar to that of a sub-lineage within another LTEE population, Ara–5, which also evolved IS*150*-mediated hypermutability, but much later in that experiment (Tenaillon et al. 2016). Most clones from the DM0 and DM25 treatments also have more deletions than the LTEE clones, again despite the fewer generations. These differences suggest some genomic instability in our study populations, in addition to the high rates of IS*150* transposition.

### Changes in growth parameters after 2,500 generation in DM0 and DM25 environments

To assess changes in growth parameters during the experiment, we compared the growth curves of evolved populations and clones to those of their respective ancestors (Supplementary Figs. S1–S10). To quantify changes in growth parameters more precisely, we estimated the slope of the log-transformed growth curves over two separate intervals in DM25. In this medium, the Cit^+^ bacteria undergo an apparent diauxic shift from growth on glucose to growth on citrate. Therefore, we chose intervals of optical density (OD) in which the change in OD over time would correspond to the respective growth rates on those resources. We also estimated the duration of the lag prior to the initial growth on glucose. A schematic of this method is shown in Supplementary Figure S1. We calibrated the relevant intervals based on the growth kinetics of two Cit^−^ strains in DM25: the founding LTEE strain, REL606 (Supplementary Fig. S2), and the anomalous evolved clone, ZDBp874 (Supplementary Figs. S5, S6). In DM0, we estimated the duration of the lag phase as well as the growth rate on citrate only.

On balance, we see the evolution of higher exponential growth rates and shorter lag phases in both DM0 and DM25; these demographic changes are consistent with those observed in the LTEE (Vasi et al. 1994). Indeed, all DM0-evolved whole populations show improvements in their growth on citrate in both DM0 and DM25 (Fig. 2), whereas their growth on glucose in DM25 shows little or no change. In DM0, the populations also exhibit markedly reduced lags prior to commencing growth (Fig. 2, Supplementary Figs. S3, S4).

**Figure 2.**
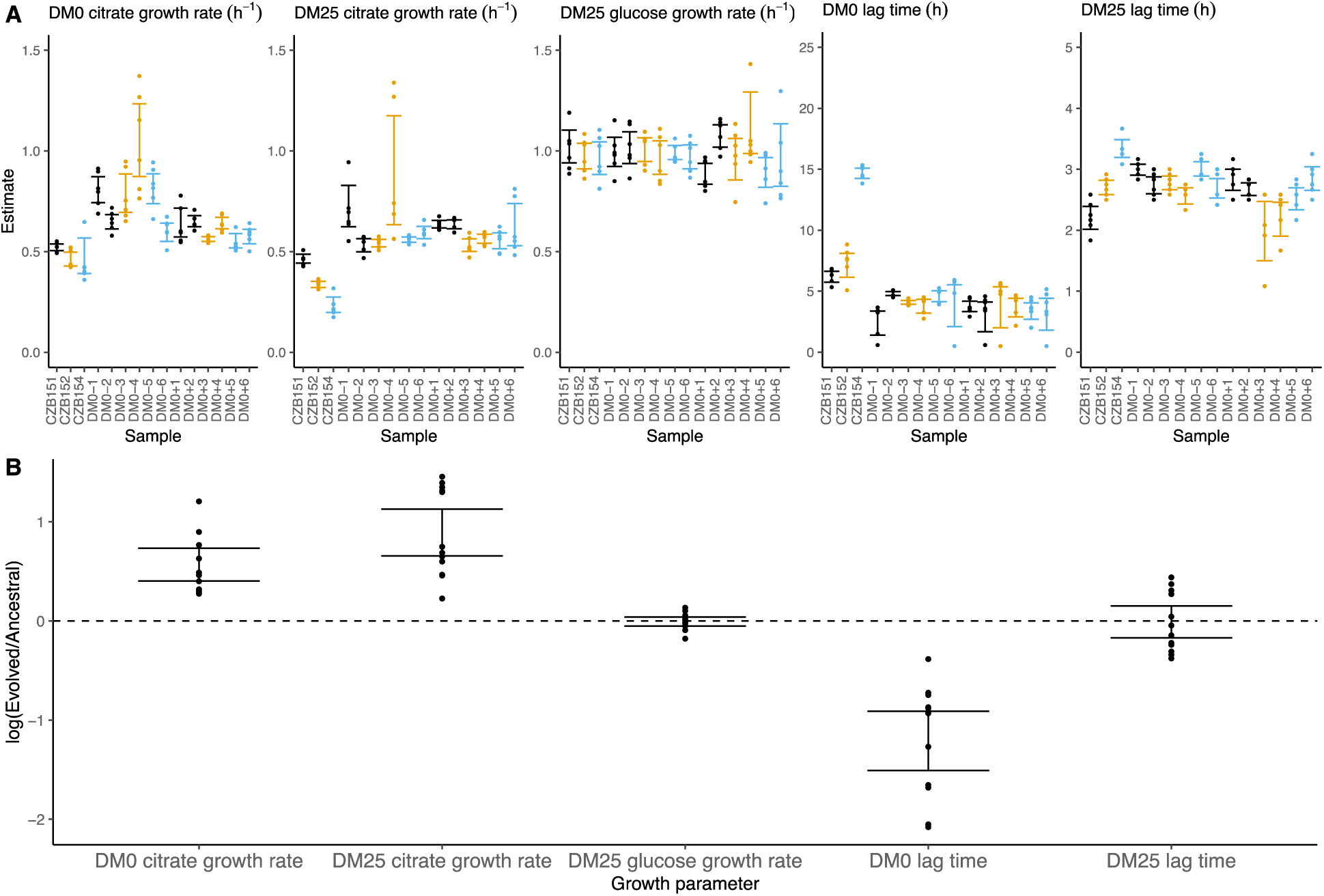
Growth parameters for whole-population samples that evolved in DM0 and their Cit^+^ ancestors. A) Estimates of various growth parameters for the ancestral strains and DM0-evolved populations at 2,500 generations, using the log-slope method. Ancestral strain CZB151 and its descendants are shown in black, CZB152 and its descendants are in orange, and CZB154 and its descendants are in black. Units for growth rates are h^−1^, and units for lag times are h. Bias-corrected and accelerated (*BC_a_*) bootstrap 95% confidence intervals around parameter estimates were calculated using 10,000 bootstraps. B) Estimates of log_2_-transformed ratios of growth parameters for the evolved populations and their ancestors. The growth curves used to estimate these parameters are shown in Supplementary Figures S3 and S4.

We observed substantially more variation in growth parameters among the evolved clones (Fig. 3, Supplementary Figs. S5–S9) than we saw when measuring the growth of whole populations. In fact, some evolved clones grow more poorly than their ancestors. Two CZB151-derived, DM0-evolved clones, ZDBp871 and ZDBp889, show little or no improvement in DM0, and they are markedly worse than CZB151 in DM25. Similarly, the anomalous Cit^−^ clone, ZDBp874, is not only unable to grow in DM0, but also grows much more poorly than its ancestor in DM25 (Fig. S5). All other DM0-evolved clones grow better than their ancestors in DM0, and most also grow about as well as their ancestors in DM25, with the additional exception of ZDBp901.

**Figure 3.**
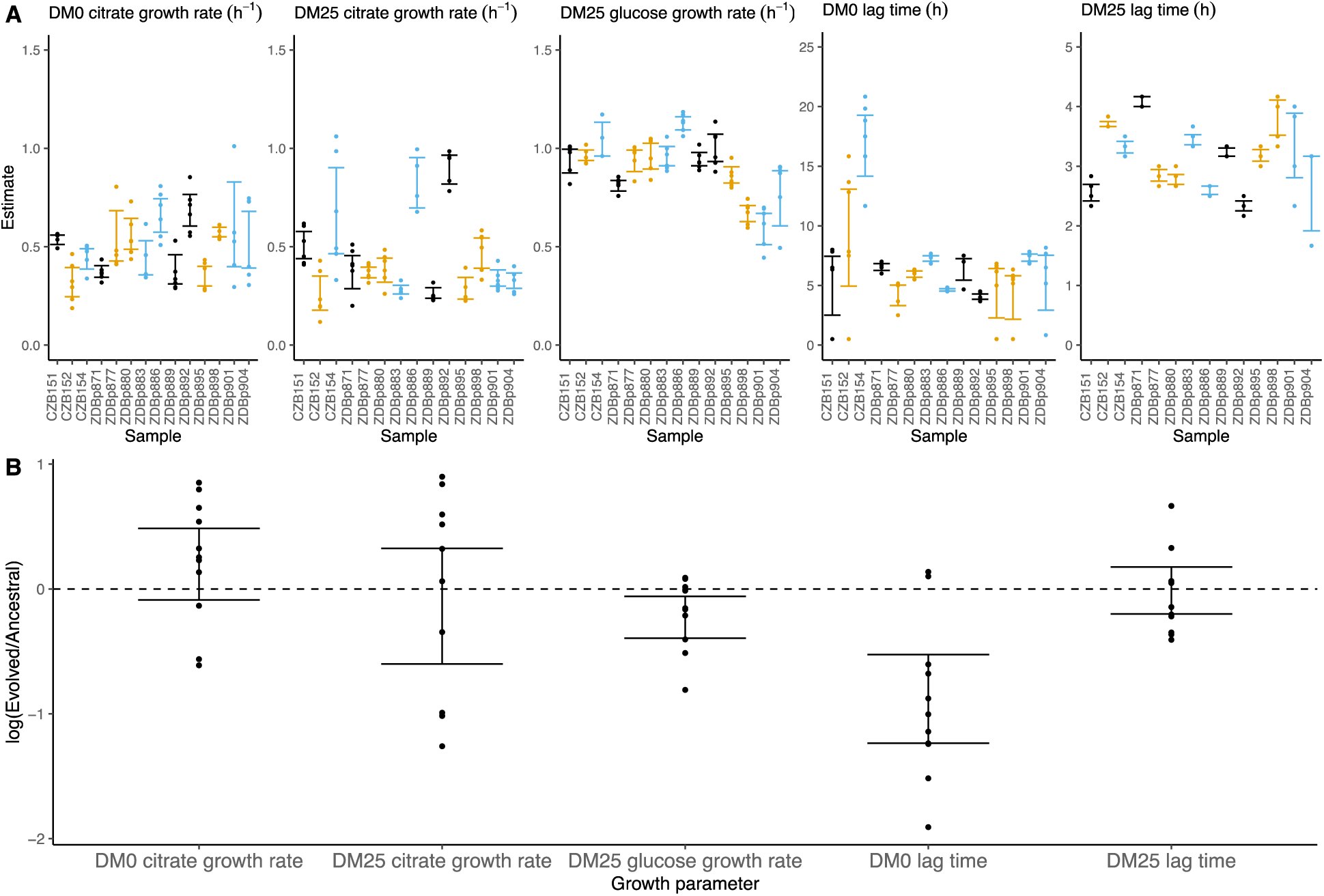
Growth parameters for clones from populations that evolved in DM0 and their Cit^+^ ancestors. A) Estimates of growth parameters for the ancestral strains and DM0-evolved clones sampled at 2,500 generations, using the log-slope method. CZB151 and its descendants are in black, CZB152 and its descendants are in orange, and CZB154 and its descendants are in blue. B) Estimates of log_2_-transformed ratios of growth parameters for the evolved clones and their ancestors. The growth curves used to estimate parameters are shown in Supplementary Figures S5 and S6; the anomalous evolved Cit^−^ clone has been excluded. See Figure 2 for additional details.

The differences in growth parameters of the evolved clones and the whole populations from which they came implies the existence of ecologically relevant genetic variation in the DM0-evolved populations. We therefore considered the possibility that improved performance on citrate might come at a cost of reduced growth on glucose. We used our separate estimates of growth rates on glucose and citrate in DM25 to determine if growth on the two substrates was correlated. However, we found no significant correlation between growth rates on citrate and glucose for either the DM0-evolved clones or whole populations (Supplementary Fig. S10A). By contrast, the growth rates measured on citrate in the two media, DM0 and DM25, are highly correlated for both clones and populations (Supplementary Fig. S10B).

### Substantial cell death in both DM0 and DM25 environments

The contribution of cell death to fitness in the LTEE is generally negligible compared to that of growth (Vasi et al. 1994). However, we serendipitously discovered evidence of substantial cell death in cultures of a Cit^+^ clone sampled from the Ara–3 population of the LTEE at 50,000 generations. We thus examined the relationship between the Cit^+^ phenotype and cell death.

We used fluorescence microscopy to quantify cell death in the DM0 and DM25 environments (Fig. 4). We analyzed five clones: the LTEE ancestor REL606; the 33,000-generation Cit^+^ CZB151; one of its DM0-evolved descendants, ZDBp871; one of its DM25-evolved descendants, ZDBp910; and the 50,000-generation Cit^+^ REL11364. We labeled cells from 24-hour stationary-phase cultures (i.e., when they would be transferred to fresh medium in the LTEE regime) using two-color live/dead stains (Methods).

**Figure 4.**
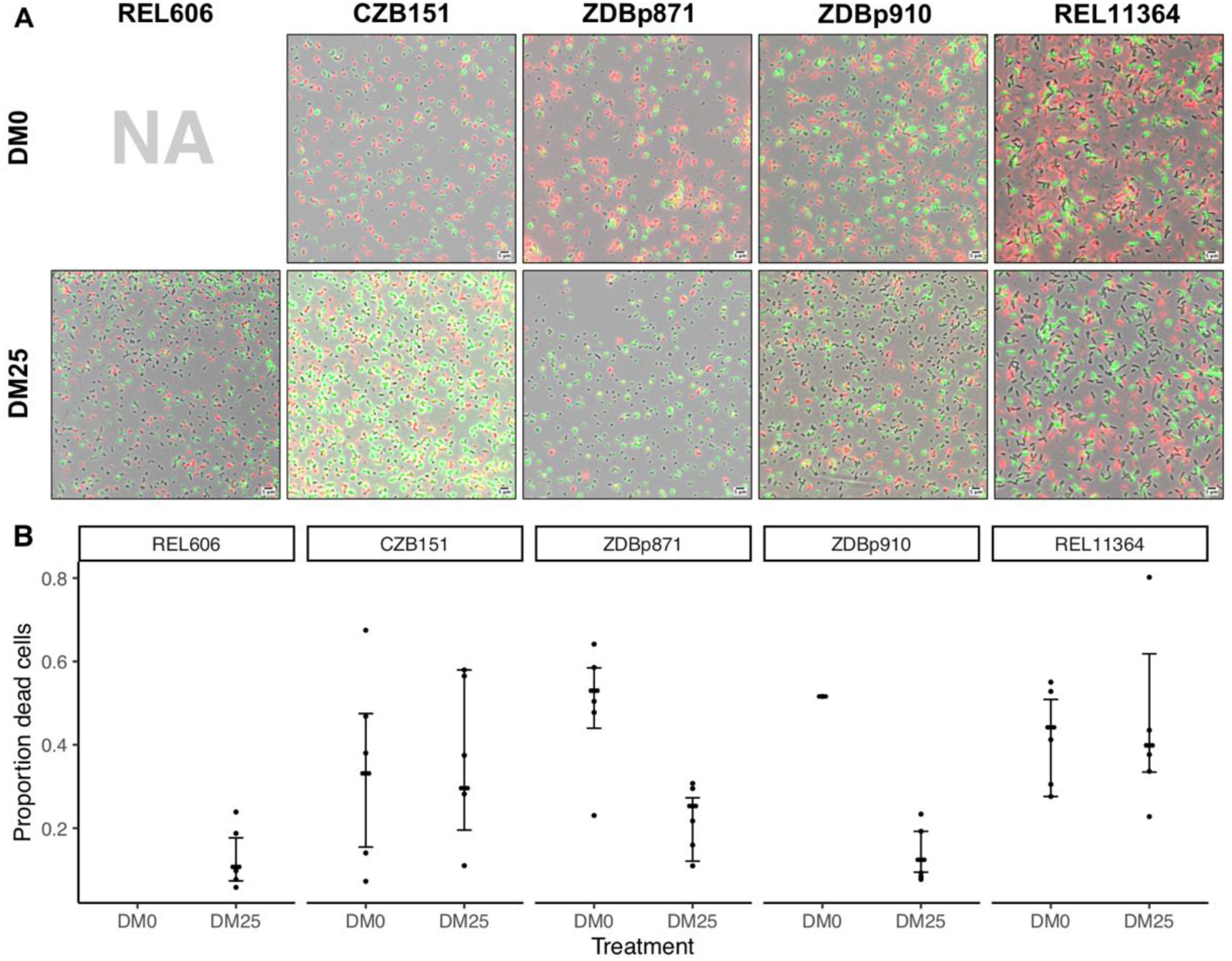
Elevated mortality in Cit^+^ strains. The Cit^+^ strains exhibit substantially elevated mortality in the citrate-only DM0 medium; some also show high mortality in DM25 as well. REL606 is Cit^−^ and cannot grow in DM0. CZB151 was sampled from LTEE population Ara–3 at generation 33,000, and its descendants ZDBp871 and ZDBp910 evolved for 2,500 generations in DM0 and DM25 media, respectively. REL11364 was sampled from LTEE population Ara–3 at generation 50,000. A) Representative micrographs of the 5 clones in the two different media. Cells were stained using the *Bac*Light Viability Kit, and they were scored as dead if their red fluorescence exceeded their green fluorescence (see Methods). Scale bars (lower right corner) represent 5 μm. B) Proportion of dead cells in 5 replicate cultures of each strain grown in DM0 and DM25 medium each (except for ZDBp910, with only one replicate). The wider symbols show estimated overall proportions weighted by the number of cells analyzed in each replicate culture. Bias-corrected and accelerated (*BC_a_*) bootstrap 95% confidence intervals were calculated using 10,000 bootstraps (except for ZDBp910), also weighted by the number of cells analyzed in each replicate.

Proportions of dead cells were calculated for 5 independent cultures for each clone and medium combination (except ZDBp910, for which we had problems with growth in DM0 and so have only one replicate, and REL606, which cannot grow in DM0). Figure 4A shows representative fields for each clone in each medium. Figure 4B shows the resulting estimates of the proportion of dead cells, along with 95% bias-corrected and accelerated (*BC_a_*) bootstrap confidence intervals (DiCiccio and Efron 1996) weighted by the number of cells analyzed and scored in the replicate cultures. On average, 10.7% of the LTEE ancestral cells grown in DM25 were scored as dead in stationary phase. By contrast, when grown in the same DM25 medium, 29.6% and 39.9% of cells were scored as dead for the Cit^+^ clones isolated from LTEE population Ara–3 at 33,000 (CZB151) and 50,000 generations (REL11364), respectively. We observed similarly high proportions of dead cells for both clones in DM0 as well (33.1% and 44.2% for CZB151 and REL11364, respectively). These results indicate that the evolution of aerobic growth on citrate was associated with elevated mortality. Moreover, the increased mortality was not remedied even after almost 20,000 generations since the new trait arose in the Ara–3 population. The two other clones we examined, ZDBp871 and ZDBp910, show somewhat different patterns. Both show lower mortality in glucose-containing DM25 (25.3% and 12.4% for ZDBp871 and ZDBp910, respectively) but higher mortality in citrate-only DM0 (53.0% and 51.6% for ZDBp871 and ZDBp910, respectively). The reduced mortality of ZDBp910 in DM25, in which it evolved for an additional 2,500 generations, suggests that the apparent metabolic imbalance associated with growth on citrate may be reduced by evolving in a medium that also contains glucose. It is even more surprising, then, that we observed no comparable reduction in mortality in the 50,000-generation Ara–3 clone, which may indicate that historically contingent ecological and genetic interactions are important for this trait. Moreover, the very high death rate of ZDBp871 in DM0, the medium in which it evolved, suggests that correcting the metabolic imbalance is even more difficult when citrate is the sole carbon and energy source.

### Specificity of genome evolution in the DM0 and DM25 environments

We found evidence that the DM0 and DM25 environments selected for mutations in different genes. Following Deatherage et al. (2017), we compared the distribution of “qualifying” mutations—nonsynonymous SNPs, deletions, duplications, and IS insertions that unambiguously affect single genes—that arose during evolution in each medium. We identified all genes in which we found at least two qualifying mutations across the 24 evolved Cit^+^ clones we sequenced. These genes are shown in Figure 5, where they are ranked by the absolute value of the difference in the number of qualifying mutations between the DM0 and DM25 conditions.

**Figure 5.**
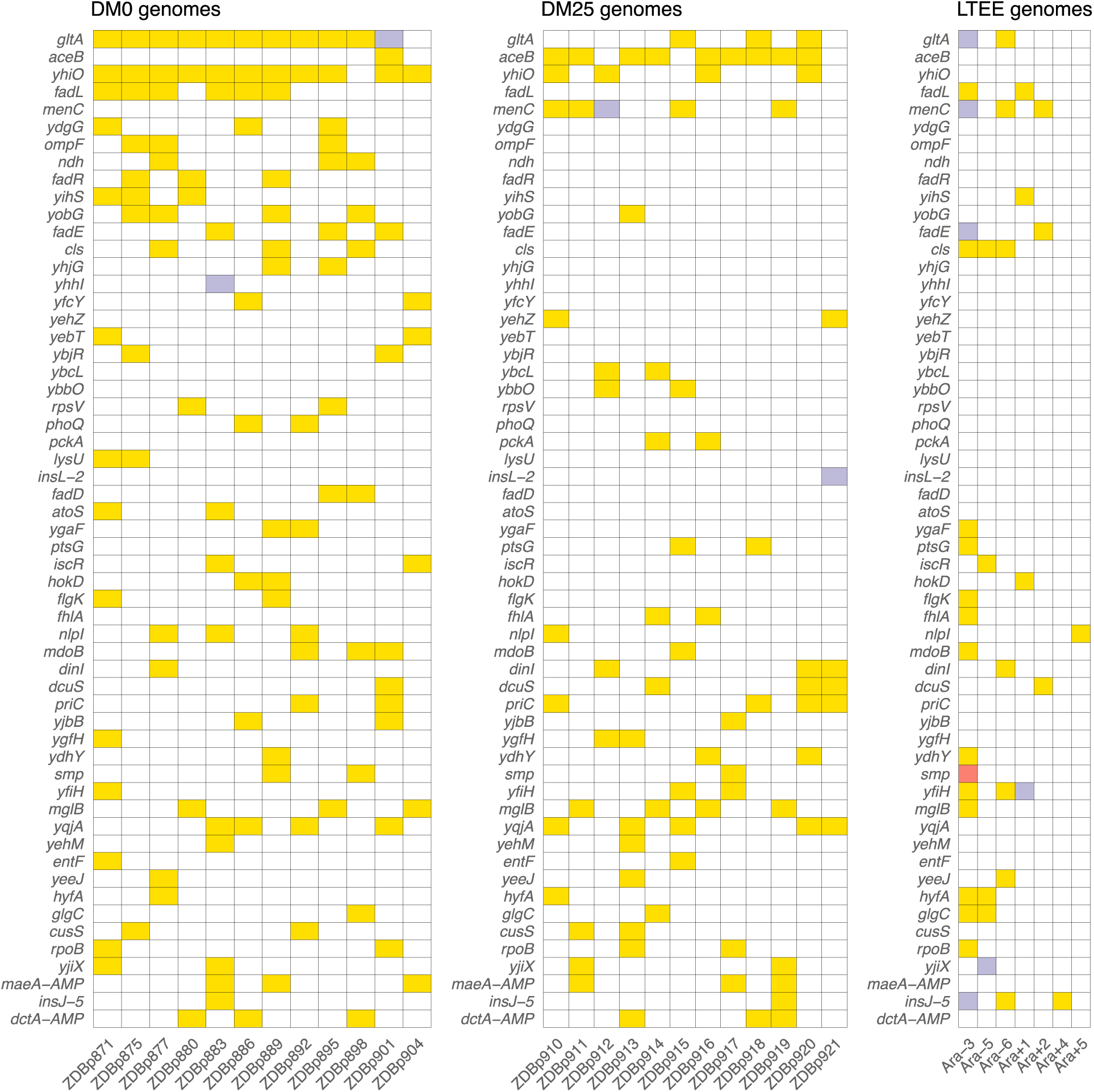
Parallel genetic evolution. Genes with mutations in two or more sequenced genomes from the DM0- and DM25-evolved populations, ranked by the absolute value of the difference in the number of qualifying mutations (see main text) between DM0 and DM25. Mutations in the same genes in the six non-mutator LTEE lineages and in a Cit^+^ clone from LTEE population Ara–3 (which evolved hypermutability), all at 50,000 generations, are shown for comparison. Yellow, violet, or red fill indicates the presence of one, two, or three qualifying mutations, respectively.

We then used the method of Deatherage et al. (2017) to quantify the extent of parallelism in genome evolution within and between the DM0 and DM25 treatments. We computed Dice’s Coefficient of Similarity, *S*, for each pair of evolved clones, where *S* = 2 |*X* ∩ *Y*|/(|*X*| + |*Y*|). |*X*| and |*Y*| are the cardinalities of the sets of genes with qualifying mutations in two clones, and |*X* ∩ *Y*| is the cardinality of the set of genes with mutations in both clones. *S* thus ranges from 0, when the two clones have no qualifying mutations in common, to 1, when both clones have qualifying mutations in exactly the same set of genes. The grand mean similarity, *S_m_*, is 0.135 across the 24 evolved clones. The mean within-treatment similarity, *S_w_*, is 0.177, meaning that two clones that evolved independently in the same medium on average have 17.7% of mutated genes in common. By contrast, the mean between-treatment similarity, *S_b_*, is 0.096, meaning that two clones that evolved in different media on average have only 9.6% of mutated genes in common. We evaluated the significance of the difference between *S_w_* and *S_b_* using a randomization test in which clones were permuted across samples 10,000 times, and the difference between the two measures was calculated for each permutation. The observed difference between the DM0- and DM25-evolved clones was higher than in any of the permutations. The greater genomic parallelism within than between environments is therefore highly significant (*p* < 0.0001).

Five genes had significantly more parallel mutations in one medium than in the other. Eleven of the 12 DM0-evolved Cit^+^ clones had qualifying mutations associated with *yhiO*, encoding the universal stress protein UspB, compared to 4 of 12 clones evolved in DM25 (Fisher’s exact test: *p* = 0.0094). Similarly, we found qualifying mutations in *gltA*, which encodes citrate synthase, in 11 of the DM0-evolved Cit^+^ clones, whereas only 3 of the DM25-evolved clones had mutations in that gene (Fisher’s exact test: *p* = 0.0028). The gene for isocitrate lyase, *aceB*, had only 1 qualifying mutation among the DM0-evolved genomes, but 9 in the DM25-evolved genomes (Fisher’s exact test: *p* = 0.0028). Among the DM0-evolved genomes, there are no qualifying mutations in *menC*, which encodes O-succinylbenzoate synthase, but 5 DM25-evolved genomes have mutations in that gene (Fisher’s exact test: *p* = 0.0373). Six of the DM0-evolved genomes have qualifying mutations in the *fadL* gene, which encodes an outer membrane long-chain fatty acid channel, but none of the DM25-evolved genomes have mutations in this gene (Fisher’s exact test: *p* = 0.0137). Moreover, we found 9 additional qualifying mutations associated with four other genes (*fadA*, *fadE*, *fadD*, and *fadR*) in the fatty-acid degradation regulon among the DM0-evolved genomes, but none in the DM25-evolved genomes. Mutations in the *fad* regulon thus show a strong signature of adaptation specific to the DM0 medium. Thirteen of the 15 total qualifying mutations in the *fad* regulon were mobile-element transpositions.

The environment was much more important than the ancestral genotype in determining the genetic targets of selection. We found no difference in the total number of qualifying mutations between the 24 evolved Cit^+^ clones when grouped by ancestor (i.e., CZB151, CZB152, CZB154) (Kruskal-Wallis test, *p* = 0.8873). Moreover, by using the same randomization test described above to test the significance of the difference between *S_w_* and *S_b_*, we found no significant difference based on ancestral genotype (*p* = 0.5540, based on 10,000 replicates).

We also found five instances of parallel changes at the amino-acid level among the DM0-evolved genomes. Three of the five occurred in *gltA*, which encodes citrate synthase: M172I, A162T, I114F. All three of these substitutions are near the allosteric binding pocket for NADH (Figure S11). Quandt et al. (2015) reported a A162V substitution that likewise affects NADH binding, and which was previously shown to fine-tune carbon flux through citrate synthase (Maurus et al. 2003). These three *gltA* mutations presumably have similar effects. We also saw parallel I197L substitutions in *ygaF*, which encodes a protein that dehydrogenates L-2-hydroxyglutarate to alpha-ketoglutarate and replenishes the cell’s reduction potential by feeding electrons from this reaction into the membrane quinone pool (Kalliri et al. 2008). There were parallel S351 substitutions in *atoS* in ZDBp871 and the anomalous Cit^−^ clone ZDBp874. This gene encodes the sensor protein of a two-component regulatory system that stimulates short-chain fatty acid catabolism. Unlike some mutations that might reduce or destroy a protein’s functionality, we expect that these parallel amino-acid substitutions fine-tune protein function (Maddamsetti et al. 2017).

### Contribution of transposable insertion elements to parallel evolution

Notwithstanding the parallel amino-acid substitutions described above, most of the parallel genomic evolution reflects the activity of IS elements. In both environments, most new IS insertions are copies of IS*150* elements (Fig. 6A, 6B). We compared the number of IS*150* insertions in clones evolved in the two media to the number that accumulated through 50,000 generations in the Ara–3 population of the LTEE (Fig. 6C). The rates of IS*150* insertion accumulation in the Ara–3 population and the DM25-evolved Cit^+^ populations are comparable, but much lower than in the DM0-evolved populations. The difference between the DM0- and DM25-evolved genomes is significant (Mann–Whitney *U* test, two-tailed *p* = 0.0089), despite the high variability between genomes within each group (Fig. 1A, 1B).

**Figure 6.**
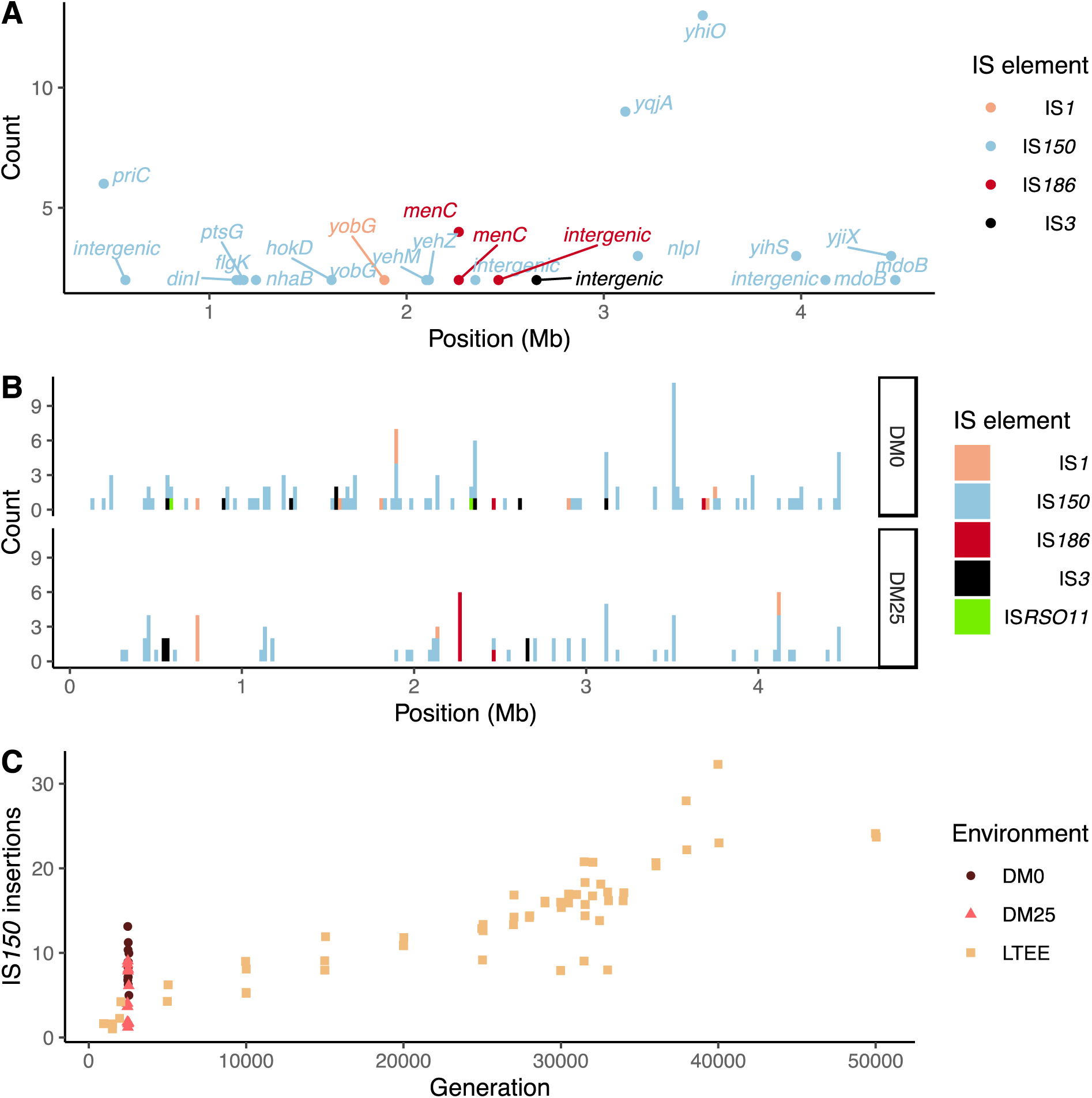
Parallel IS-element insertions. A) Counts of parallel IS-element insertions in labeled genes (including promoter and coding regions) summed across sequenced DM0- and DM25-evolved genomes, and arranged by position on the *E. coli* chromosome, relative to the inferred last common ancestor of all strains (Methods). *IS*1 insertions are shown in pink, IS*150* in light blue, IS*186* in red, IS*3* in black, and IS*RSO11* in chartreuse. Some genes contain multiple sites with parallel IS-element insertions. B) Location of insertions, shown separately for the DM0- and DM25-evolved genomes. Colors are the same as in panel A. C) Total number of IS*150* insertions in the DM0- and DM25-evolved genomes after 2,500 generations. Corresponding number in clones from LTEE population Ara–3 over 50,000 generations are shown for comparison. DM0 clones are labeled as brown circles, DM25 clones as pink triangles, and LTEE Ara–3 clones as tan squares.

Insertions of IS*150* into new sites were strongly parallel across the independently evolved populations within, but not between, the two environments (Fig. 6A, 6B). These systematic differences led us to hypothesize that the parallel IS insertions reflect the influence of selection, rather than insertion-site biases (Figs. 5, Fig. 6B) (Tenaillon et al. 2016). We evaluated this hypothesis by conducting a statistical test for selection-driven parallel IS*150* insertions over and above a null model that assumes only insertion-site preferences. In this test, we conservatively assumed that IS*150* can only insert into the positions in which we observed insertions in one or more sequenced genomes from either this experiment or the LTEE (Tenaillon et al. 2016). We also assumed that the probability of IS*150* transposing into a given site is proportional to the observed number of IS*150* insertions at that site across the sequenced genomes, as would be the case if mutational biases alone accounted for the parallel IS*150* insertions. We then used the non-parametric bootstrap method (100,000 replicates) to calculate the probability that any particular site would be hit by so many IS*150* elements among the DM0-evolved genomes, holding the number of IS insertions over that group fixed. The resulting probability that 9 or more IS*150* insertions would occur at the same site across the 12 DM0 populations is ∼0.01. The most extreme observed case of parallelism at the base-pair level was an IS*150* insertion at the –35 position of the promoter for *yhiO*, which encodes the universal stress protein UspB, which happened in 9 of the 12 DM0 genomes. This test is even more conservative because it excludes two other IS*150* insertions affecting this same gene in the DM0 genomes: an IS*150* insertion at the –36 position of the promoter and another IS*150* insertion in *yhiO* itself. Therefore, we can confidently reject the null hypothesis that site-specific insertion biases alone provide an adequate explanation for the distribution of IS*150* insertions. Given the conservative nature of this test, it is quite possible that some other parallel IS-insertions also indicate positive selection.

### Parallel amplification mutations in the DM0- and DM25-evolved populations

We detected tandem amplifications of large genome regions, often to very high-copy-number, in many of the DM0- and DM25-evolved clones (Table 1, Fig. 7, Supplementary Table S1). All genomes include amplifications containing the novel genetic module that evolved during the LTEE, which places one or more copies of the *citT* gene under the control of the *rnk* promoter region, with the exception of the anomalous Cit^−^ clone ZDBp874. This new *rnk-citT* module provides access to citrate, and mutations that increase its dosage improve growth on citrate (Blount et al. 2012, Van Hofwegen et al. 2016).

**Figure 7.**
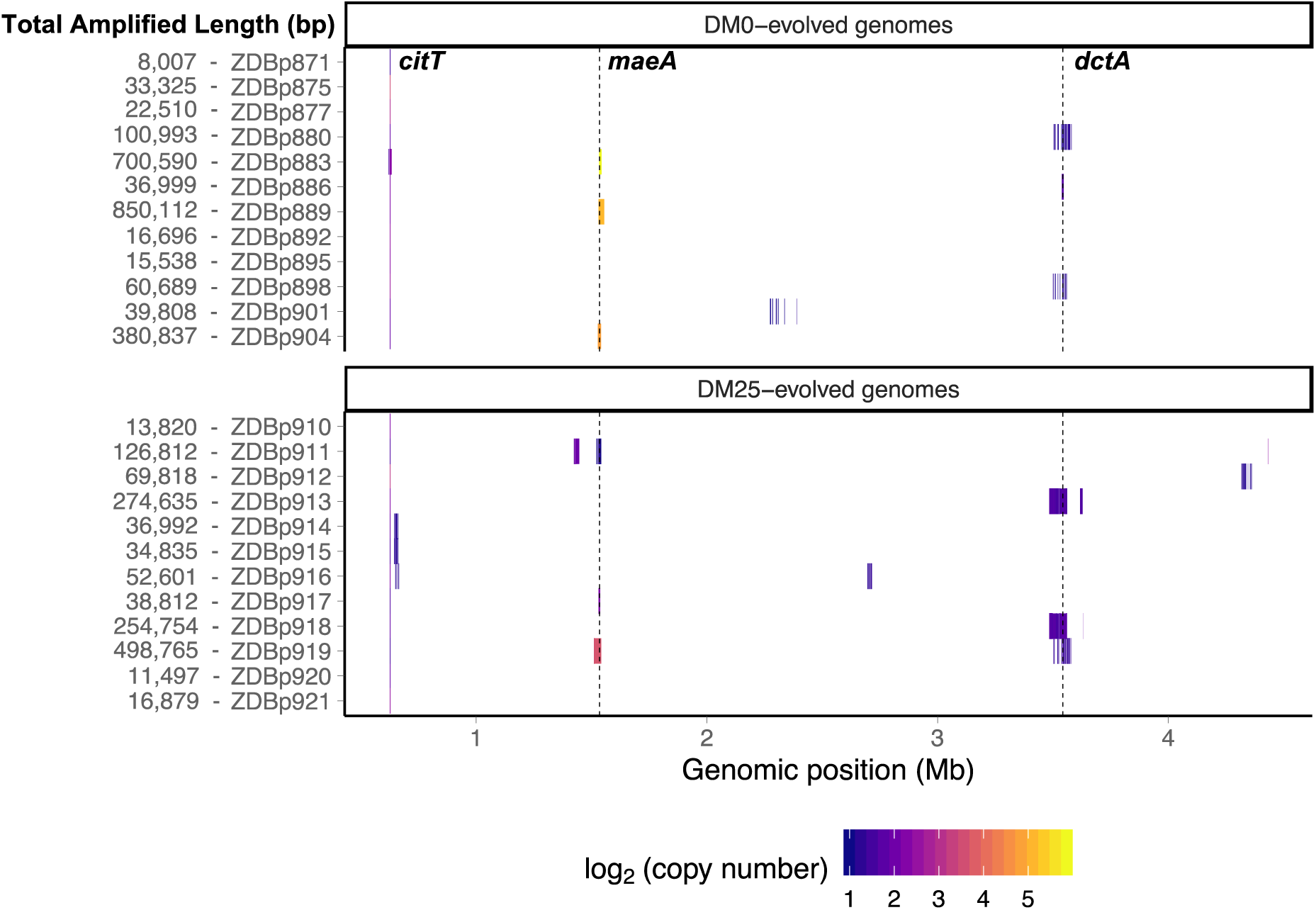
Genetic amplifications in evolved clones. A) Genomic regions with significant amplifications in DM0- and DM25-evolved clones, arranged by chromosomal position. The evolved clones from DM0 (top half) and DM25 (bottom half) are indicated at the near left, with the total amplified length shown at the far left. Dashed vertical lines mark the *maeA* and *dctA* loci. The boundaries vary among the subset of genomes with amplifications that encompass these genes; by contrast, the *citT* locus is amplified in all of these genomes, and with nearly uniform boundaries. Colors denote amplification copy-number on a log_2_ scale from dark (low copy-number) to light (high copy-number).

**Table 1:** Copy number of amplified *maeA* and *dctA* genes in sequenced clones from populations that evolved for 2,500 in either DM0 or DM25 environments.

| Genome | Medium | Gene | Mean copy number | Minimum number* | Maximum number* | Adjusted *p*-value** |
| --- | --- | --- | --- | --- | --- | --- |
| ZDBp880 | DM0 | *dctA* | 2.33 | 1.67 | 3.26 | <0.0001 |
| ZDBp886 | DM0 | *dctA* | 3.20 | 1.88 | 4.25 | <0.0001 |
| ZDBp898 | DM0 | *dctA* *** | 2.09 | 1.71 | 2.76 | 0.0023 |
| ZDBp913 | DM25 | *dctA* | 3.17 | 1.91 | 4.68 | <0.0001 |
| ZDBp918 | DM25 | *dctA* | 3.41 | 1.80 | 5.19 | <0.0001 |
| ZDBp919 | DM25 | *dctA* | 2.60 | 2.06 | 3.29 | <0.0001 |
| ZDBp883 | DM0 | *maeA* | 58.47 | 22.61 | 95.46 | <0.0001 |
| ZDBp889 | DM0 | *maeA* | 34.71 | 2.35 | 55.39 | <0.0001 |
| ZDBp904 | DM0 | *maeA* | 28.08 | 15.30 | 44.72 | <0.0001 |
| ZDBp911 | DM25 | *maeA* | 2.22 | 1.54 | 3.39 | <0.0001 |
| ZDBp917 | DM25 | *maeA* | 4.72 | 3.56 | 6.18 | <0.0001 |
| ZDBp919 | DM25 | *maeA* | 12.81 | 2.16 | 18.00 | <0.0001 |

\*These bounds indicate the ratio of the minimum and maximum sequencing coverage measured at the indicated locus to the mean sequencing coverage over the genome.
\*\*Significance levels are shown after Bonferroni corrections for multiple tests of the same hypothesis.
\*\*\*There may be two discontinuous amplifications of *dctA* in this genome, or there may be a single continuous amplification with a short region of low coverage within the gene. The second region of amplification has a similar copy number. We present data for only one region, which provides a conservative estimate of the overall statistical significance in this case.

Other amplifications include the *dctA* gene. DctA is a proton motive force-driven, generalized di- and tricarboxylic acid transporter. During growth on citrate, the CitT antiporter protein exports TCA cycle intermediates into the medium in exchange for citrate. DctA permits recovery of those intermediates. Two mechanisms of increasing *dctA* expression have been shown to improve growth on citrate. Quandt et al. (2014) identified mutations in the *dctA* promoter that cause constitutive, high-level expression. Van Hofwegen et al. (2016) showed that increased dosage of *dctA* is likewise beneficial. We found evidence that these two mechanisms are anticorrelated. Two of the ancestral clones, CZB151 and CZB154, have a shared (identical by descent) mutation in the promoter sequence of *dctA*. The third ancestor, CZB152, lacks this mutation. Only one of the 16 evolved descendants of CZB151 or CZB154 has a *dctA* amplification, whereas five of CZB152’s eight descendants have such an amplification (Fisher’s exact test: *p* = 0.0069). Also supporting this anticorrelation, one of the three CZB152 descendants without a *dctA* amplification independently evolved a mutation affecting that gene’s promoter.

We identified another set of parallel amplifications in six evolved genomes. These amplifications are large and highly variable in extent, though all include at least the *fdnI*, *yddM*, *adhP, maeA, rpsV,* and *bdm* genes. The amplifications were often present in high copy numbers. Three DM25-evolved genomes have 2-13 copies, and three that evolved in DM0 have 28-59 copies (Table 1). In one case, ZDBp889, the amount of DNA in the amplified region constitutes more than 15% of the total evolved genome (Fig. 7). By contrast, the amplifications of *citT* and *dctA* have average copy numbers of 4-5 and 2-3 copies, respectively (Table 1, Supplementary Table S1).

Given their considerable length and high copy number, these amplifications must exert a metabolic burden owing to the additional DNA and increased gene expression (DaSilva and Bailey 1986; Lenski and Nguyen 1988). The fact that amplifications of this genomic region evolved repeatedly suggests that they also confer some selective benefit that outweighs their cost. We examined the genes shared among the amplifications to identify which might confer this benefit. The *rpsV* gene, which encodes the 30S ribosomal subunit protein D, appears to have been a minor target for adaptation to DM0 based on parallel mutations (Fig. 5). The *maeA* gene encodes an NAD^+^-dependent oxaloacetate-decarboxylating malate dehydrogenase (EC 1.1.1.38) that catalyzes the decarboxylation of malate to pyruvate. This plausible connection to citrate metabolism led us to hypothesize that increased *maeA* dosage and expression provides the benefit that overcomes the cost imposed by the amplifications.

### Increased MaeA expression is highly beneficial in the citrate-only environment

We tested our hypothesis that increased *maeA* dosage confers a fitness benefit by transforming the CZB151 and CZB152 ancestral strains with a low-copy plasmid, RM4.6.2, which contains a copy of *maeA* that is under the control of a strong constitutive synthetic promoter and ribosome-binding site. These Ara^−^ RM4.6.2 transformants were competed in DM0 against Ara^+^ mutants (ZDB67 and ZDB68, respectively) of the same clones transformed with the empty-plasmid control. The RM4.6.2 plasmid conferred a ∼27% selective advantage in both the CZB151 (*n* = 6; mean fitness = 1.2790, *t*-distributed 95% confidence interval: [1.2636, 1.2944]) and CZB152 (*n* = 6; mean fitness = 1.2778, *t*-distributed 95% confidence interval: [1.2597, 1.2959]) backgrounds relative to their otherwise isogenic competitors. Overexpression of *maeA* is therefore highly beneficial during growth in DM0, and its benefit likely explains the large, high-copy-number amplifications containing *maeA* found in many evolved clones.

We used RNA-Seq to verify that clones with *maeA*-containing amplifications have elevated transcription of that gene. We compared the transcriptomes of two ancestral clones (CZB151 and CZB152) and two DM0-evolved clones with *maeA* amplifications (ZDBp883, ZDBp889). Both evolved clones do, indeed, have much higher levels of *maeA* expression than their respective ancestors (Supplementary Fig. S12).

Despite the large fitness advantage conferred by increased *maeA* dosage in the citrate-only DM0 environment, most of the evolved Cit^+^ genomes we examined do not have *maeA* amplifications. Moreover, whereas three of the DM25-evolved genomes in this study have large *maeA* amplifications (Table 1), none have been found in the sequenced genomes of any Cit^+^ clones isolated from the Ara–3 parent population in the LTEE itself. This discrepancy might be explained by the evolution of increased *maeA* expression via other mutations. To evaluate this possibility, we also used RNA-Seq to examine the transcriptome of ZDBp877, a DM0-evolved clone without a *maeA* amplification. In contrast to ZDBp883 and ZDBp889, ZDBp877 expresses *maeA* at a level similar to that of the ancestral clones (Supplementary Fig. S12). This finding means that at least some, and perhaps all, of the evolved clones without *maeA* amplifications lack mutations that boost its expression. In any case, the mechanistic basis for the fitness benefit conferred by *maeA* remains unknown.

### Transcriptomic analysis of DM0-evolved clones

We identified other potentially adaptive differences in transcription between the DM0-evolved clones during growth in the DM0 medium (Supplementary Figure S12). The two evolved clones with *maeA* amplifications, ZDBp883 and ZDBp889, both show increased expression of the *fad* fatty acid β-oxidation regulon, whereas ZDBp877, which lacks a *maeA* amplification, does not. ZDBp877 and ZDBp889, but not ZDBp883, both downregulate the cytochrome *bo_3_* terminal oxidase complex, *cyoABCD*. We also found three genes with more extreme differential expression than *maeA* between the clones with and without the *maeA* amplification. These genes are *dinI, gltS, and ECB_03510*, all three of which are strongly downregulated in ZDBp877 in comparison to both clones with the amplification. DinI is a DNA-damage inducible protein that regulates the SOS response. GltS is a glutamate/sodium symporter. *ECB_03510* encodes a protein of unknown function, and it lies immediately downstream of *gltS*.

These differences aside, we found largely similar changes in gene expression across the three DM0-evolved clones relative to their ancestors (Supplementary Figure S12). All three evolved clones show strong downregulation of the UspB stress protein encoded by *yhiO*, presumably caused by the parallel IS*150* insertions into that gene’s promoter. We also found extensive downregulation of genes encoding ribosomal proteins (including *rpsB*, *rpsU*, *rpsO*, *rpsT, rplE, rplJ*, *rplN*, and *rplX*); genes involved in RNA transcription (*rpoA, rpoB, rpoC, rpoS*, *rho*); and DNA-replication associated genes (*gyrA*). Other down-regulated genes in the evolved clones include the *nuo* operon, which encodes NADH dehydrogenase in the respiratory electron transport chain, and key TCA cycle genes including those encoding the 2-oxoglutarate dehydrogenase complex (*sucB*, *sucC*, and *lpdA*). By contrast, we see strong upregulation of genes encoding certain prophage-associated proteins (ECB_00826 and ECB_00827); some toxin-antitoxin pairs (*chpA*-*chpR*); proteins involved in recombinational DNA repair (*recA* and *recN*); SOS response proteins (*dinD*, *sulA*, *umuC*, and *umuD*); proteins associated with stationary phase (*csiE* and *sbmC*); a biofilm-associated stress protein (*bhsA*); and others of unknown significance (Supplementary Files S2 and S3).

The downregulation of transcription, translation, and NADH dehydrogenase genes together with the increased expression of stress-associated genes suggests that adaptation to DM0 might have involved reducing growth rate, presumably to achieve balanced growth on citrate alone. The upregulation of fatty-acid β-oxidation genes (albeit less so in the ZDBp877 clone without the *maeA* amplification) also indicates some remodeling of the connection between fatty acid and citrate metabolism that is mediated by acetyl-CoA. Other changes, including the upregulation of the *fad* operon encoding fatty acid degradation in ZDBp883 and ZDBp889; the anaerobic glycerol-3-phosphate dehydrogenase operon (*glpABCD*) in ZDBp877 and ZDBp889; and the glycerol-3-phosphate transporter (*glpD* and *glpT*) in ZDBp883 (Supplementary File S3) suggest adaptation to scavenging on dead and dying cells in the DM0 populations.

## DISCUSSION

It is rarely feasible to examine evolution in action as organisms invade and colonize a new niche in natural environments, especially with independently evolving replicates and control populations. In this study, we investigated how *E. coli* variants with the novel ability to grow aerobically on citrate adapted to a novel, citrate-only resource environment in the laboratory. We examined the genomic and phenotypic evolution of 12 initially clonal populations after 2,500 generations in this new environment, along with 12 initially identical control populations maintained for the same time in the ancestral environment that contains glucose as well as citrate, to better understand their post-invasion potential, including refinement of the Cit^+^ trait.

### Growth and death in the novel environment

The founding clones grew poorly in their new medium, with extended lag phases, slow growth, and high variation in their growth kinetics. However, the founding clones clearly possessed substantial potential to adapt to citrate as their sole carbon and energy source, as they evolved shorter lag times and faster growth rates in DM0. They also showed correlated improvements in the ancestral glucose-citrate medium, DM25. These changes are consistent with selection pressures typical for evolution experiments that use a serial batch-culture regime like that of the LTEE (Vasi et al. 1994), upon which our experiment was based.

In contrast to the pervasive improvements observed at the population-level, we found substantial variation in the growth phenotypes of the evolved clones. While most showed large improvements similar to what we saw at the population level, some evolved clones grow only slightly better, or even worse, than their ancestors. For example, one clone from a control population that evolved in DM25, ZDBp917, had a lag time far longer than its ancestor. Such paradoxical phenotypic evolution could represent the sampling of standing deleterious variation that has not yet been purged by selection. This possibility is consistent with the high rate of IS transposition observed in these populations. Perhaps more likely, the clones with paradoxical phenotypes could be ecotypes adapted to unknown niches, such as scavenging on dead cells or cross-feeding on metabolites produced by other, co-existing lineages (Turner and Souza et al. 1996, Rozen et al. 2009, Velicer and Mendes-Soares 2009, Le Gac et al. 2012, Maddamsetti et al. 2015, Good et al. 2017). In that case, the paradoxical phenotypes of some clones might hint at the evolution of ecological complexity in these populations. We will investigate the ecological dimension of these populations’ evolution in future work.

We found substantial numbers of dead cells in cultures of both DM0- and DM25-evolved clones as well as a Cit^+^ clone isolated at generation 50,000 of the LTEE (Figure 4). This finding is puzzling because all of these clones were sampled from populations that had experienced long periods of adaptation to growth on citrate. Theory also predicts that mutations that reduce the mortality of daughter cells should be highly favored when rare (Wahl and Dai Zhu 2015). A number of hypotheses might explain the unexpected cell death. Cell death may be a direct physiological consequence of the Cit^+^ trait; for example, death might result from unbalanced growth on citrate. Alternately, death might be a side effect of Cit^+^ refinement. Under either scenario, cell death could persist if the benefit of growing on citrate exceeds the cost of elevated mortality. Still another alternative explanation would be mechanistic constraints on Cit^+^ refinement that cause antagonistic pleiotropy; for example, adaptations that allow faster growth or shorter lags on citrate may be worth the cost of increased death. High death rates would then be a correlated response to selection for improved growth on citrate that could persist indefinitely, or until compensatory mutations arise that break the pleiotropic tradeoff. Finally, the ancestral clones might have constrained potential to refine the Cit^+^ trait, such that the supply of beneficial mutations that decrease cell death when growing on citrate is limited. Although later evolution might increase the supply of viability-refining mutations, the maladaptive mismatch between cell physiology and the new ecological niche could persist for a long time.

Several facts weigh against the side-effect and weak-selection hypotheses as being sufficient to explain our results. We found that cell death remains high in DM0 after 2,500 generations of growth on citrate, and in DM25 nearly 20,000 generations after the Cit^+^ trait arose in the Ara–3 population of the LTEE. The difference in death rates observed in DM0 and DM25 indicates that growth in citrate alone remained especially stressful for some cell lineages, even after thousands of generations of adaptation. Moreover, cell death releases resources that could be scavenged by other cells. Any genetic tendency toward death might exert a double fitness penalty in that it would both decrease that lineage’s net growth while increasing that of competing scavenger lineages in these well-mixed and spatially unstructured conditions. Thus, antagonistic pleiotropy seems the most likely explanation. Adaptations that confer a large enough growth advantage to outweigh any pleiotropic costs associated with death would be successful in a novel niche.

### Genome evolution during adaptation to the new environment

Comparing the ancestral and evolved genomes allowed us to identify a variety of genomic changes underlying the phenotypic adaptation that occurred in our experiment. The pace of genomic evolution was faster, in both our experimental and control populations, than in the LTEE populations that did not evolve hypermutable phenotypes. Genome evolution was also faster in the citrate-only DM0 environment than in DM25. The elevated rate of SNP accumulation might be due in part to the additional cell divisions that can occur in bacterial cultures that experience significant death (Frenoy and Bonhoeffer 2018). However, much of the increased rate of genome evolution reflects elevated IS element activity; IS*150* copies were particularly active. Stress de-represses transposase expression and activity, resulting in bursts of transposition and IS-mediated mutagenesis (Miousse et al. 2015, Vandecraen et al 2017, Fouché et al. 2019). The high levels of IS activity are thus consistent with other indications that growth on citrate is stressful, and that growth in a citrate-only resource environment is particularly so. Differences in the nature and intensity of selection between different environments, and even between different populations experiencing the same environment, will also influence the spread of mutations, especially beneficial ones, making it difficult to fully understand the observed trends.

The IS-mediated mutations we identified had a variety of effects. Some IS copies inserted into genes, likely eliminating their function. Others occurred in intergenic regions, where they may have altered gene expression. We identified several parallel insertions that occurred independently in multiple populations, suggesting that they were beneficial. For instance, parallel insertions of IS*150* in the promoter of the *yhiO* gene that encodes universal stress protein B (UspB) were present in most of the DM0-evolved clones. Our transcriptomic data indicate that these insertions repress UspB expression. Overexpression of UspB has been shown to cause increased cell death in stationary phase (Farewell et al. 1998), and so these insertions might be related to the elevated cell death we observed. More generally, our findings are consistent with work showing that IS and other transposable elements are rich sources of genetic variation and genomic plasticity during adaptive evolution, particularly given bursts of transposition caused by host stress (Casacuberta and Gonzaléz 2013, Wright et al. 2017, Dubin et al 2018, Fouché et al 2019, Sentausa et al. 2019).

The most dramatic genomic changes we found were amplifications, some of which were quite large and high in copy number. These included amplifications containing *citT*, which encodes the antiporter protein that provides access to citrate, and *dctA*, which permits recovery of C_4_-dicarobxylates exported into the medium during growth on citrate. Increased dosages of both genes have been shown to be beneficial for growth on citrate, so these amplifications are not unexpected (Blount et al 2012, Quandt et al. 2015). We also identified large amplifications containing *maeA*, which in one evolved clone comprised 15% of its total genome size. We showed that overexpression of *maeA* from a plasmid confers a substantial fitness benefit in DM0. We suspect that this benefit comes from the resulting effect on carbon flow through biosynthetic pathways, and we hope to test this hypothesis in future work.

Although the *maeA* amplifications confer a large fitness advantage, only a minority of the evolved clones in our study possess such amplifications. Unlike the case of *dctA*, we did not identify mutations that were anti-correlated with the *maeA* amplifications, nor did we find any mutations in the *maeA* promoter indicative of cis-regulatory changes. Clones lacking these amplifications also showed no increased expression of *maeA*. Other mutations may have occurred in the other populations that make *maeA* overexpression unnecessary, although we do not know which mutations might do so. We also cannot exclude the possibility that the *maeA* amplifications might be common in other populations, but their size and copy number make them prone to collapse by homologous recombination. In that case, evolved clones lacking amplifications may have lost them during the course of their isolation. Finally, it is also possible that clones without *maeA* amplifications are simply less fit than those with the amplifications, leaving the door open for later amplifications if our evolution experiment had run longer.

### Conclusions

Our findings show that genomic plasticity due to transposable-element activity and copy-number variation can be an important contributor to post-innovation adaptation to new niches. These findings support and extend previous work showing the importance of such plasticity in adaptation to other selective challenges (Chang et al. 2013, Vandecraen et al. 2017, Press et al. 2019, Lauer et al. 2019), including the rapid evolution of antibiotic heteroresistance in clinical pathogens (Nicoloff et al. 2019). In particular, our work bolsters earlier demonstrations of the selective value of dynamic gene amplifications that ameliorate metabolic bottlenecks by increasing the dosage of genes encoding specific products needed at higher levels (Patrick et al. 2007, Andersson and Hughes 2009). For instance, Blank and colleagues (2014) found that *E. coli* strains with single-gene knockouts rapidly re-evolved the capacity to grow in minimal medium in part via amplifications that increased genome size by more than 20%. Such amplifications can carry substantial metabolic costs and are prone to recombination-mediated collapse, so they are readily lost when the relevant gene products are no longer valuable. However, amplifications also increase the opportunity for more stable mutations that may stabilize the benefit on a single copy, favoring subsequent collapse and elimination of the cost of the high copy number (Andersson and Hughes 2009, Brennan et al 2015). With more time to adapt, the *maeA* amplifications found in some of the evolved populations might thus be supplanted by more stable SNP or indel mutations that increase expression without the metabolic costs of the amplifications.

We also found that transposon activity contributed to faster genomic evolution in our experimental and control populations than has been typical in most LTEE populations (among those that did not evolve point-mutation hypermutability). IS-mediated insertional inactivation of genes has been shown to contribute to virulence (Hammerschmidt et al. 1996), improved growth on glucose (Gaffé et al. 2011), and resistance to various antibiotics (Hernandez-Alles et al. 1999, Fujimura and Murakami 2008). IS insertions can also alter the regulation of genes either by providing outward-directed promoters or by the fortuitous creation of new promoters (Mahillon and Chandler 1998). Transposable element-mediated gene regulation changes have been shown to alter host range (Coleman et al. 2014), enable growth on citrate (Blount et al. 2012), and confer antibiotic resistance (Sóki et al. 2013). IS elements seem to have both inactivated and modulated the expression of various genes in our evolution experiment. IS activity is increased by stress, which cells might experience when they invade a new niche to which they are poorly adapted (Vandecraen et al. 2017). Of course, this plasticity is a double-edged sword: the genomic instability that transposable elements cause can also produce deleterious mutations that could impede adaptation and might even lead to extinction, especially when small founding populations invade a new niche.

While our findings support the consensus understanding of the evolutionary effects of amplifications and IS activity, it is difficult to disentangle and quantify the contributions of mutation and selection to these genome dynamics. Little is currently understood about the molecular basis for the substantial variation in the rates and site-specific biases of IS transposition across bacterial strains, species, IS elements, and environments even in the absence of natural selection (Nzabarushimana and Tang 2018). Rigorous analysis of larger datasets will be needed to develop the necessary general approach and models.

Despite the evidence for adaptation in our experimental populations, the high rates of cell death we observed make clear that the evolved cells remain imperfectly adapted. The physiological cause of the high death rates remains unclear, as does the reason it persists despite substantial time for its amelioration. Further work will be needed to discriminate among the various hypotheses that might explain the ultimate causes of such persistent mortality: weak selection to overcome it, antagonistic pleiotropy (tradeoffs with other important traits), limited genetic potential for refinement, or some combination thereof. Further work is also necessary to identify the proximate mechanisms causing cell death in this system.

Altogether, our results suggest that genomes often possess a latent potential for adaptation to new niches following an innovation-driven discovery. This potential can be fulfilled not only by point mutations, but also via amplification mutations and IS element activity that may allow more rapid adaptation after a novel trait provides access to a new niche. Provided that there is no competition from resident populations or other invasive lineages, and provided that the niche is stable, then access to the new niche may virtually guarantee successful establishment. However, the persistent high rates of cell death we observed suggest that long periods of adaptation may sometimes be necessary before a new ecotype becomes fully fitted to its new position in nature.

Going forward, our experimental system has the potential to address questions about innovation, adaptation, and maladaptation that are relevant to both evolutionary biology and evolutionary medicine. Evolutionary innovations can result in maladaptive mismatches between physiology and environment, similar to those we have seen in our experimental populations. Among humans, for example, cultural innovations, including the agricultural revolution, reshaped both diets and gut microbiota. Contemporary high-calorie diets and sedentary lifestyles have caused an epidemic of metabolic, diabetic, and obesity-related illnesses. How common is this pattern in evolution? Do evolutionary innovations generally result in metabolic and physiological problems? Might new traits and others that have been recently reshaped by evolution have elevated phenotypic variation, lower robustness, or both? If growth on citrate in *E. coli* requires major changes in physiology and metabolism, then that innovation may have increased fragility as a consequence of difficulties in coordinating cell growth and division with the substantially remodeled physiology and metabolism (Scott et al. 2014, Schaechter 2015). By disrupting existing physiological and metabolic processes, new traits may produce altered selection pressures that require novel variation to solve the resulting imbalances. That new variation may, in turn, adversely affect correlated traits and the organism’s overall robustness. We conjecture that the refinement of evolutionary innovations that provide organisms with exclusive access to new niches may, in general, tilt the balance between evolvability and physiological robustness in living systems toward evolvability (Lenski et al. 2006), at the expense of robustness and overall good health.

## METHODS

### Evolution experiment

Three Cit^+^ clones were previously isolated at random from the 33,000-generation sample of LTEE population Ara–3, and designated as CZB151, CZB152, and CZB154 (Blount et al. 2008). Spontaneous Ara^+^ revertants were isolated for each clone and designated as ZDB67, ZDB68, and ZDB69, respectively. Isolated colonies of each clone and its revertant were inoculated into Luria Bertani (LB) broth, grown overnight at 37°C with orbital shaking at 120 rpm, and frozen at –80°C with 0.8% glycerol as cryoprotectant for long-term preservation. The clones and revertants were later revived from the frozen stocks in LB and grown overnight. The revived cultures were then preconditioned by 10,000-fold dilution into 9.9 mL Davis Mingioli (DM) minimal medium supplemented with 25 mg/L glucose (DM25) and grown for 24 h at 37°C with orbital shaking. Cultures were then diluted 100-fold into 9.9 mL of fresh DM25 and grown for another 24 h. This preconditioning acclimated the bacteria to growing on citrate. The preconditioned cultures were then diluted 100-fold into 9.9 mL DM basal medium (DM0), in which 1 g/L (1,700 mM) citrate is the sole carbon source. Two replicate populations were started for each LTEE-derived clone and revertant, for 12 total DM0 populations. At the same time, 12 populations were inoculated into DM25. The DM25 populations were maintained at 37°C with orbital shaking and transferred by 100-fold dilution into fresh DM25 every 24 h (i.e., the same conditions as in the LTEE) for 375 transfers and 2,500 generations in total. The founding Cit^+^ clones grow poorly in the citrate-only resource environment, and thus were unable to reach stationary phase or, in some cases, exponential phase within 24 or even 48 h. The DM0 populations were therefore incubated for 72 h after initial inoculation to permit them to reach stationary phase before transfer to fresh medium. They were then diluted 100-fold into 9.9 mL of DM0 every 48 h for seven cycles (two weeks), and then subsequently every 24 h for a total of 375 transfers and 2,500 generations. Every 37 days (∼250 generations) samples of each population were frozen with glycerol at –80°C.

### Isolation of evolved clones

Each evolved population sample was revived by inoculating 100 μL of the stock frozen at generation 2,500 into 9.9 mL of LB broth and incubated overnight at 37°C with 120 rpm orbital shaking. The revived DM0- and DM25-evolved populations were then diluted 10,000-fold in 9.9 mL of DM0 or DM25, respectively, grown for 24 h at 37°C with orbital shaking, followed by 100-fold dilution in fresh DM0 or DM25 and another 24 h of growth at 37°C with orbital shaking. Each population was then diluted 100,000-fold in 0.85% saline and spread on an LB agar plate marked with 3 dots on the bottom. The plates were incubated at 37°C for 48 h, after which the colony closest to each dot was streaked for isolation on an LB plate, thereby providing three randomly chosen clones from each population. An isolated colony of each clone was then inoculated into LB broth, grown overnight, and frozen as before.

### Growth curves

One of the three evolved clones was chosen from each DM0 or DM25 population, revived, and preconditioned in DM0 or DM25 as described above. The cultures were then diluted 100-fold into 9.9 mL of DM0 or DM25, vortexed, and six 200-μL aliquots of each culture dispensed into wells in a 96-well plate. Optical density (OD) at 420-nm wavelength was measured every 10 min for 48 h using a Molecular Devices SpectraMax 384 automated plate reader. Measurements before the 30-minute-mark were discarded from the analysis.

### Microscopy and cell viability analyses

We performed microscopy and viability analyses on cells derived from five clones: the LTEE ancestor (REL606); one of the three Cit^+^ ancestors in our evolution experiment (CZB151); two of its descendants that evolved in DM0 and DM25 for 2,500 generations (ZDBp871 and ZDBp910, respectively); and a Cit^+^ clone isolated at generation 50,000 of the LTEE (REL11364). We revived clones from the frozen stocks and preconditioned them as described above, except that the preconditioning steps in DM0 or DM25 were extended to 4 daily passages to ensure acclimation to these environments. Preparations for live/dead cell staining and microscopic analyses were performed on the fifth day. We concentrated the cells in each culture by centrifugation at 7,745 g for 8 min and decanted the supernatant. Cell pellets were resuspended in Corning tubes containing 10 mL of 0.85% saline and incubated at room temperature for 1 h; the tubes were inverted every 15 min. We then centrifuged these cultures for an additional 8 min, decanted the supernatant, and resuspended the cell pellets in 0.85% saline. The volume of saline was adjusted for variation in turbidity, in order to ensure that we had sufficient cells in a typical field of view for microscopy. We examined 14-55 fields per replicate for each combination of strain and media treatment. Total cell counts ranged from approximately 15,000 to 60,000 for the various combinations of clones and culture media.

We used the LIVE/DEAD *Bac*Light Viability Kit for microscopy (ThermoFisher #L7007), following the manufacturer’s directions for fluorescently labeling cells. In short, we mixed components A and B in equal amounts, added 1 µl to each culture containing resuspended cells, and incubated them for 20 min in the dark to prevent photobleaching. After labeling, we fixed 3 µL of each sample onto a 1% agarose pad and performed fluorescent microscopy using a Nikon Eclipse Ti inverted microscope. Phase-contrast images were taken using diascopic light with an exposure time of 100 ms. Fluorescence was measured with an exposure time of 200 ms at 25% power of the fluorescent light source using two filter sets, 49003-ET-EYFP and 49008-ET-mCherry Texas Red (Chroma), which correspond to the fluorescence spectra of “live” and “dead” cells, respectively. All images were taken at 100× magnification.

Micrographs were analyzed using *SuperSegger*, an image-processing package (Stylianidou et al. 2016). We first filtered the data, keeping only those values for segmented regions in the micrograph that were scored by the neural-network classifier as having *P*(Cell = True) > 75%. (Region scores range between –50 and 50, so we used data only from regions with values between 25 and 50). We then used the fluorescence values from the *SuperSegger* output and scored individual cells as “live” or “dead” depending on whether the fluorescence signal on the green (YFP) channel was greater or lesser, respectively, than the signal on the red (RFP) channel. We then calculated the proportion of dead cells across the many fields examined for each of the 5 replicate cultures that we analyzed for each combination of clone and growth medium, and we used these values in the statistical analyses.

### Genomic analysis and copy-number variation

The 3 Cit^+^ founder strains (CZB151, CZB152, CZB154), their respective Ara^−^ derivatives (ZDB67, ZDB68, ZDB69), and 25 evolved clones (one Cit^+^ clone from each DM0 and DM25 evolved population, plus the anomalous Cit^−^ clone ZDBp874) were thawed and grown overnight in LB medium. Genomic DNA was isolated from each sample using the Qiagen Genomic-tip 100/G DNA extraction kit. The genomic DNA was then sequenced by the facilities and using the platforms shown in Supplementary File S4.

For genomes sequenced at UT Austin, DNA was purified from *E. coli* cultures using the PureLink Genomic DNA Mini Kit (Invitrogen). For each sample, 1 µg of purified DNA was fragmented using dsDNA Fragmentase (New England Biolabs). Then, the KAPA Low Throughput Library Preparation kit (Roche) was used to construct Illumina sequencing libraries according to the manufacturer’s instructions with two exceptions. First, reaction volumes were cut in half. Second, DNA adapters were designed that incorporate additional 6-base sample-specific barcodes such that the barcodes are sequenced as the first bases of both read 1 and read 2. Paired-end sequencing with 300-base reads was performed on an Illumina MiSeq at the University of Texas at Austin Genome Sequencing and Analysis Facility. Reads were demultiplexed using a custom python script. Barcodes and adapter sequences were trimmed using *Trimmomatic* version 0.38 (Bolger et al., 2014).

When available, short-read data from different sequencing platforms were combined before mutation identification. We identified mutations in the sequenced genomes using *breseq* version 0.33.2 (Deatherage and Barrick 2014). A bash script called “generate-LCA.sh” was used to infer the last common ancestor (LCA) of all evolved strains by taking the intersection of mutations found in previously curated genomes for CZB152 and CZB154; those curated founder genomes (and others) are available at: https://github.com/barricklab/LTEE-Ecoli. The mutations called by *breseq* relative to the LCA were further analyzed using custom python and R scripts that are available, and described more fully, at: https://github.com/rohanmaddamsetti/DM0-evolution.

The following algorithm was used to find copy-number variation in the genomes. The *breseq* pipeline models 1× copy number with a negative binomial distribution fit to coverage, truncating high and low coverage that might be caused by amplifications and deletions, respectively. We then identified all positions in the genome that rejected that negative binomial at an uncorrected *p* = 0.05. Finally, we calculated a Bonferroni-corrected *p*-value for contiguous stretches of the genome in which the 1× null model was rejected at each site. We examined coverage at sites separated by the maximum sequencing read length to ensure they were not spanned by a single read. For example, in the case of a region of elevated coverage that was 1000 bp in length, covered by 150-base Illumina sequencing reads, the value of *P*(*coverage*=*min*)^6^ would be calculated, where *min* is the minimum coverage in that region, *P*(*coverage*=*min*) is the probability of that minimum coverage under the negative binomial null model, and 6 represents the (integer) number of sites that are 150 bp apart in the 1000-bp stretch. The output was then filtered for regions longer than 2 × 150 = 300 bp to remove potential false positives. The Bonferroni calculation included corrections for checking every site in the genome in addition to the number of sites that passed the initial 0.05 cutoff for deviations from the negative binomial expectation. All gene amplifications detected in the DM0- and DM25-evolved genomes are reported in Supplementary File S1.

### RNA-Seq and transcriptome analysis

We performed RNA-Seq on six clones: the three Cit^+^ clones from the LTEE used as ancestors in our evolution experiment (CZB151, CZB152, and CZB154) and three evolved descendants isolated after 2,500 generations of adaptation to DM0 (ZDBp877, ZDBp883, and ZDBp889). Each clone was revived from a frozen stock in LB as described above. The cultures were diluted 10,000-fold into DM25 with four-fold replication and grown for 24 h at 37°C with 120 rpm orbital shaking for preconditioning to minimal medium. The 16 resulting cultures were then diluted 100-fold in DM0 and grown for 48 h at 37°C with shaking for preconditioning to the citrate-only medium. The cultures were then diluted 100-fold again into fresh DM0, and grown to OD_600_ 0.2 – 0.3, corresponding to mid-log phase, at which point their RNA was extracted using the cold phenol-ethanol method (Bhagwat et al. 2003). RNA was recovered using a Qiagen RNeasy MiniKit (#74104), and DNA was removed with a Qiagen RNase-free DNase set (#79254). RNA was diluted to 50 ng/mL with nuclease-free water and cDNA amplified by RT-PCR. Purified cDNA was then sequenced by Admera Health (South Plainfield, NJ). We used *kallisto* version 0.44 (Bray et al. 2016) to quantify RNA transcripts and *sleuth* (Pimentel et al. 2017) to conduct differential-expression analysis and visualization. These results are presented in Supplementary File S2.

### Construction of maeA plasmid

We constructed a medium-copy-number plasmid based on the kanamycin resistance cassette-containing plasmid, pSB3K3, in which the *maeA* gene was placed under the control of a strong constitutive synthetic promoter and ribosome binding site, P089-R052, described by Kosuri et al. (2013). We used PCR to amplify the *maeA* gene from REL606 and the pSB3K3 plasmid. We ordered the P089-R052 promoter as an oligonucleotide. We assembled these components using circular polymerase cloning (Quan and Tian 2009) and Gibson assembly (Gibson 2011). Drop dialysis using Millipore membrane filters (VSWP01300) was performed for 15 min to desalt the assembly reactions before electroporation. We isolated transformants on LB-Kanamycin plates and used PCR to find colonies that contained the P089-R052–*maeA* insert. We used Sanger-sequencing of plasmid inserts to verify that no unintended point mutations had occurred during construction. The final plasmid containing the P089-R052-*maeA* insert in the pSB3K3 backbone was named RM4.6.2.

### Competition experiments to assess fitness effects of maeA

The Cit^+^ ancestral clones CZB151 and CZB152 and their Ara^+^ revertants, ZDB67 and ZDB68, respectively, were transformed with the plasmid RM4.6.2. We also transformed the same clones with the empty pSB3K3 vector. Stock cultures of each transformant were then frozen at –80°C with glycerol as a cryoprotectant.

Each RM4.6.2 transformant competed against its cognate pSB3K3 transformant in the clone with the opposite Ara marker state. Briefly, all 8 transformants were revived in LB supplemented with 50 μg/mL kanamycin and grown overnight at 37°C with 120 rpm orbital shaking. Each overnight culture was then diluted 10,000-fold in 9.9 mL DM0 and incubated for 48 h at 37°C with orbital shaking, after which it was diluted 100-fold in fresh DM0 every 48 h three times to acclimate cells to the citrate-only resource environment. Competition assays commenced the next day by inoculating 9.9 mL DM0 with 50 μL each of an RM4.6.2 transformant and the oppositely marked pSB3K3 transformant, with 4-fold replication for a total of 16 competitions. Three-day competitions were then conducted as described in detail elsewhere (Lenski et al. 1991).

## Data availability statement

All analysis and statistical scripts will be deposited at www.datadryad.org (DOI: XXXX) upon acceptance. RNA-Seq data have been deposited in the NCBI SRA under accession PRJNA553503. Genome sequencing data have been deposited in the NCBI SRA under accession PRJNA595472. Analysis code is also available at: https://github.com/rohanmaddamsetti/DM0-evolution.

## Acknowledgments

We thank Joshua Franklin and Yann Dufour for helpful discussions and assistance with the microscopy work; Simon D’Alton for assistance with genome sequencing; Daniel Barich for help in handling transcriptomics data; Jessica Baxter and Neerja Hajela for assistance in the laboratory; Jean Vila, Erik Quandt, Daniel Deatherage, Dacia Leon, Debora Marks, David Ding, Yarden Katz, Helen Murphy, and Kyle Card for helpful discussions; and Sandeep Venkataram and Sébastien Wielgoss for useful feedback on an early version of the manuscript.

## SUPPLEMENTARY INFORMATION

**Supplementary File S1.** Full results for gene amplifications in the DM0- and DM25-evolved genomes.

**Supplementary File S2.** Full results of differential expression analysis between ancestral (CZB151 and CZB152) and evolved strains (ZDBp877, ZDBp883, ZDBp889), calculated using the Wald test implemented in *sleuth* (Pimentel *et al*. 2017).

**Supplementary File S3.** Details of populations, clones, and genome sequencing datasets described in this study.

**Supplementary Table S1.** Copy number of amplified *citT* genes in sequenced clones.

| **Genome** | **Medium** | **Mean copy number** | **Minimum copy number\*** | **Maximum copy number\*** |
| --- | --- | --- | --- | --- |
| CZB151 | DM25 | 4.21 | 3.39 | 5.47 |
| CZB152 | DM25 | 8.23 | 5.17 | 11.46 |
| CZB154 | DM25 | 4.14 | 1.72 | 9.83 |
| ZDBp871 | DM0 | 2.82 | 1.70 | 4.26 |
| ZDBp875 | DM0 | 11.37 | 8.05 | 14.93 |
| ZDBp877 | DM0 | 7.68 | 3.66 | 11.33 |
| ZDBp880 | DM0 | 3.82 | 1.79 | 5.93 |
| ZDBp883 | DM0 | 5.08 | 1.88 | 12.81 |
| ZDBp886 | DM0 | 4.76 | 2.27 | 6.88 |
| ZDBp889 | DM0 | 4.66 | 2.90 | 7.11 |
| ZDBp892 | DM0 | 5.69 | 2.21 | 8.76 |
| ZDBp895 | DM0 | 5.30 | 2.13 | 8.51 |
| ZDBp898 | DM0 | 6.14 | 2.77 | 9.73 |
| ZDBp901 | DM0 | 3.91 | 1.83 | 5.64 |
| ZDBp904 | DM0 | 3.84 | 1.78 | 5.68 |
| ZDBp910 | DM25 | 4.71 | 2.40 | 6.47 |
| ZDBp911 | DM25 | 3.13 | 1.58 | 5.01 |
| ZDBp912 | DM25 | 8.93 | 4.51 | 13.41 |
| ZDBp913 | DM25 | 4.83 | 2.68 | 6.94 |
| ZDBp914 | DM25 | 4.11 | 2.03 | 6.17 |
| ZDBp915 | DM25 | 3.31 | 1.78 | 5.19 |
| ZDBp916 | DM25 | 4.87 | 2.67 | 6.95 |
| ZDBp917 | DM25 | 3.20 | 1.77 | 4.63 |
| ZDBp918 | DM25 | 3.66 | 1.88 | 5.18 |
| ZDBp919 | DM25 | 2.91 | 2.06 | 3.81 |
| ZDBp920 | DM25 | 3.92 | 2.08 | 5.49 |
| ZDBp921 | DM25 | 5.76 | 2.93 | 9.15 |

\*These bounds indicate the ratio of the minimum and maximum sequencing coverage measured at the *citT* locus to the mean coverage over the genome. In all cases, the estimated copy number is significantly greater than 1 at $p < 0.0001$ , even after Bonferroni corrections for multiple tests of the same hypothesis.

**Figure S1.**
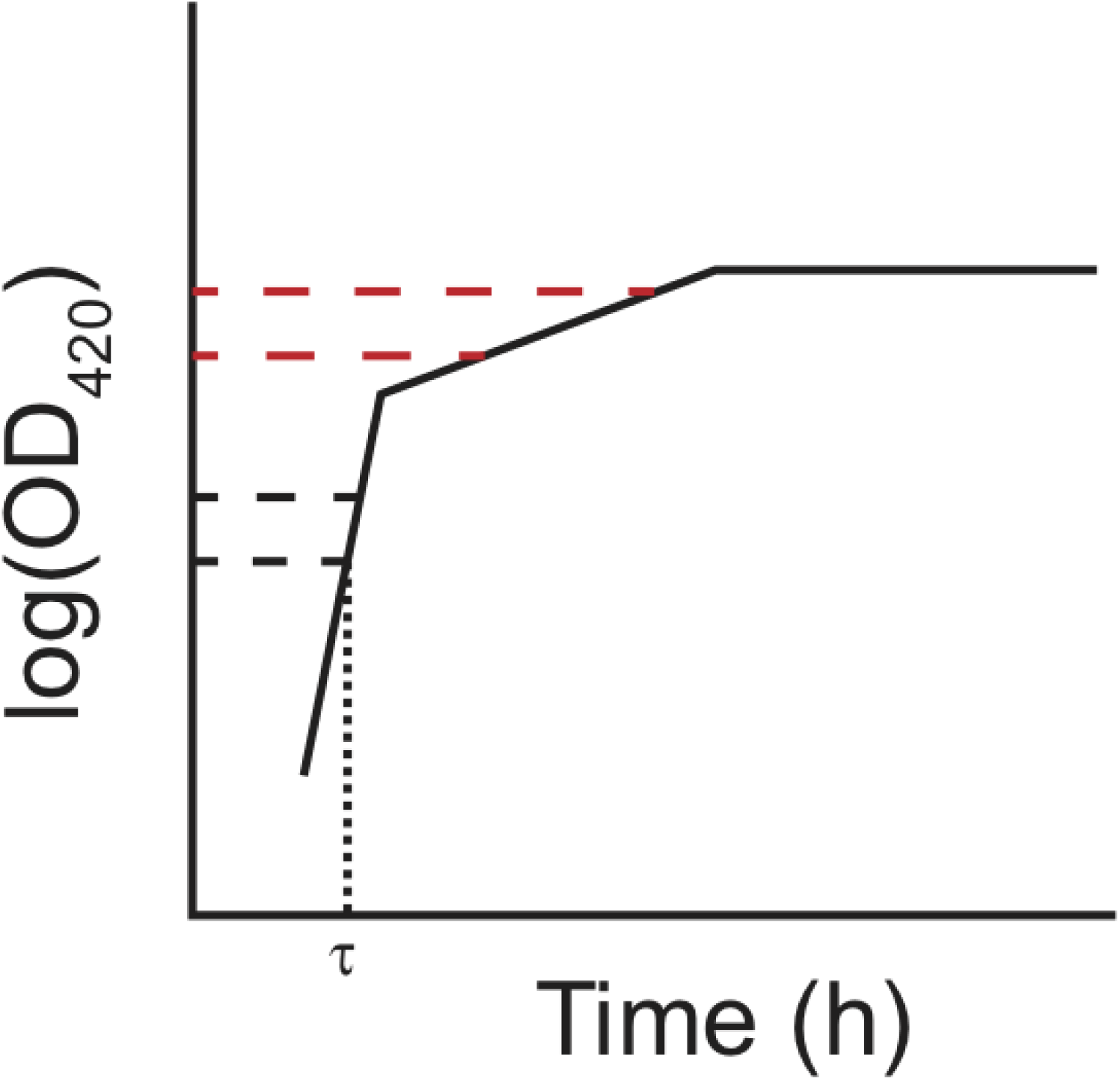
Schematic of the log-slope method to calculate growth rates. Optical densities were log*_e_*-transformed, and the slope of the curve in the interval OD_420 nm_ = [0.01, 0.02] was used to calculate the exponential growth rate on glucose (h^−1^), *r_glucose_*. The slope of the curve in the interval OD_420 nm_ = [0.05, 0.1] was used to calculate the exponential growth rate on citrate (h^−1^) *r_citrate_*. This interpretation assumes a diauxic shift between growth on glucose and citrate, rather than simultaneous utilization of both substrates. In any case, growth rates during these intervals are relevant phenotypes even without assuming diauxie. A lag time (τ) was estimated as the time (h) until OD_420 nm_ = 0.01 was reached.

**Figure S2.**
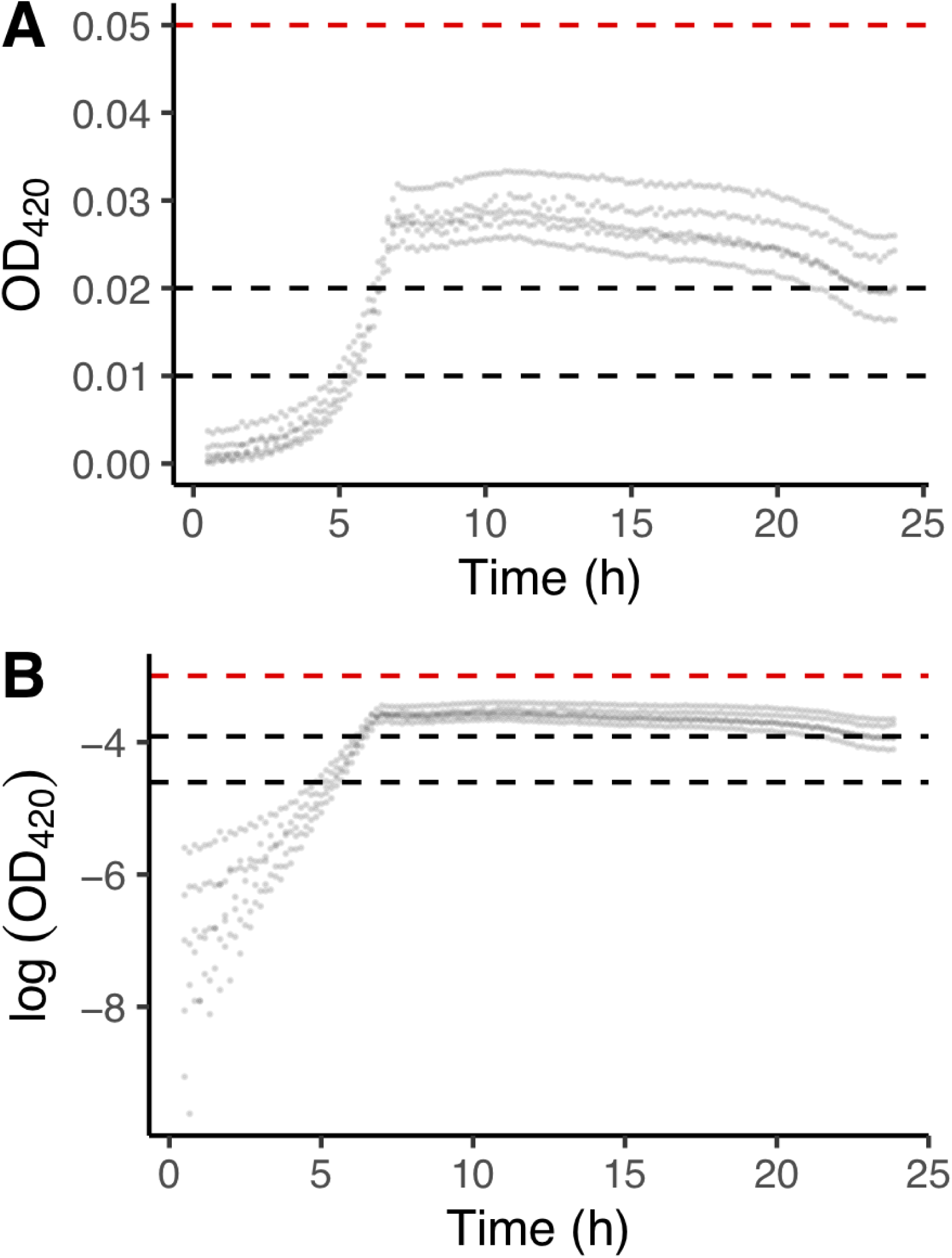
Growth curves for REL606 in DM25. These data were used to choose the interval for estimating the exponential growth rate on glucose. A) Replicated growth curves in DM25. B) The same data as in panel A except log*_e_*-transformed. Dashed black lines indicate the interval used to calculate growth rates on glucose; the dashed red line shows the lower bound of the interval in which the growth rate on citrate would be estimated (Supplementary Fig. S1).

**Figure S3.**
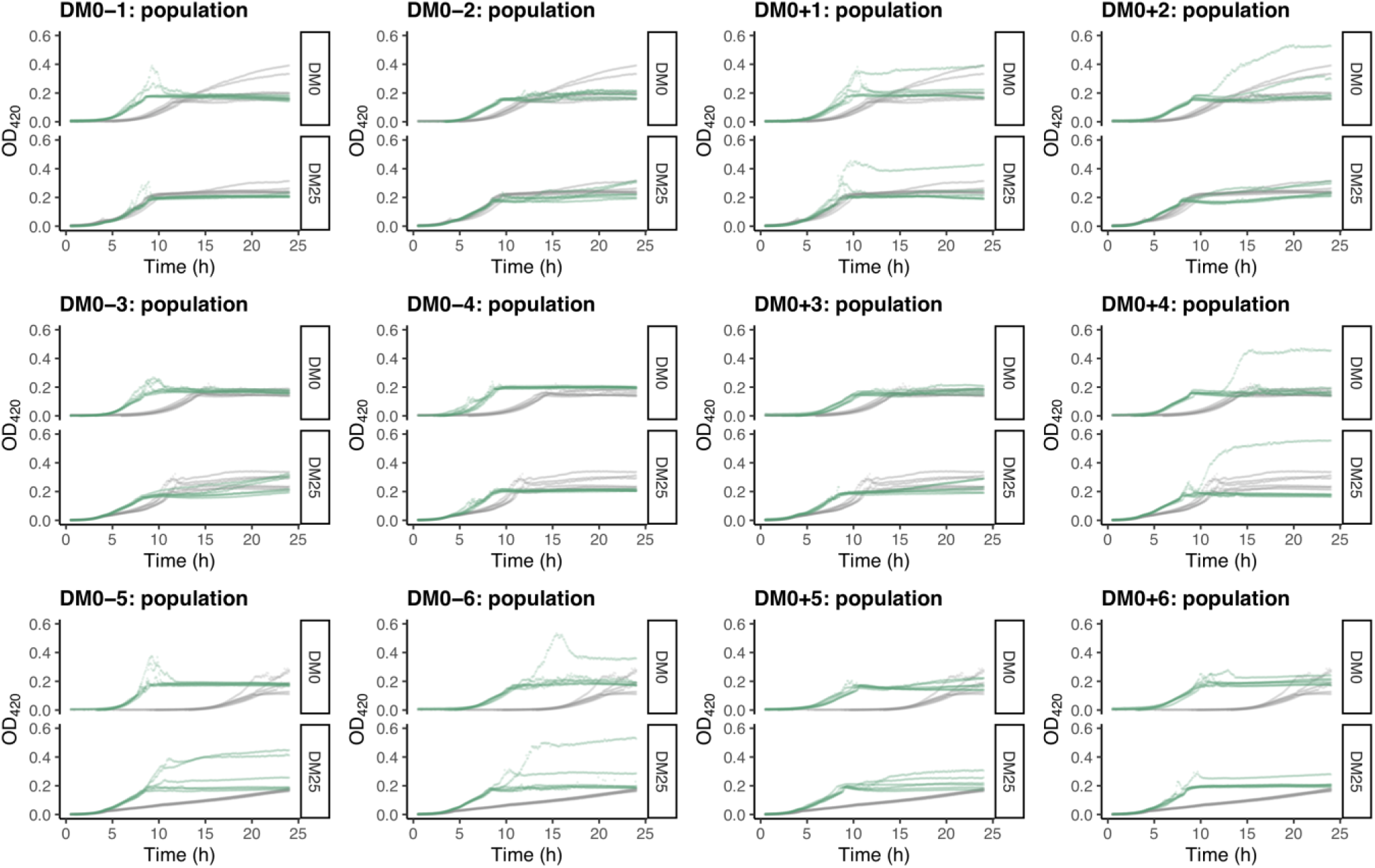
Growth curves of the 12 DM0-evolved whole-population samples, measured in DM0 and DM25. For comparison, growth curves of the evolved populations are paired with those of their ancestors: CZB151 (top row), CZB152 (middle row), and CZB154 (bottom row). The evolved and ancestral curves are shown in green and gray, respectively.

**Figure S4.**
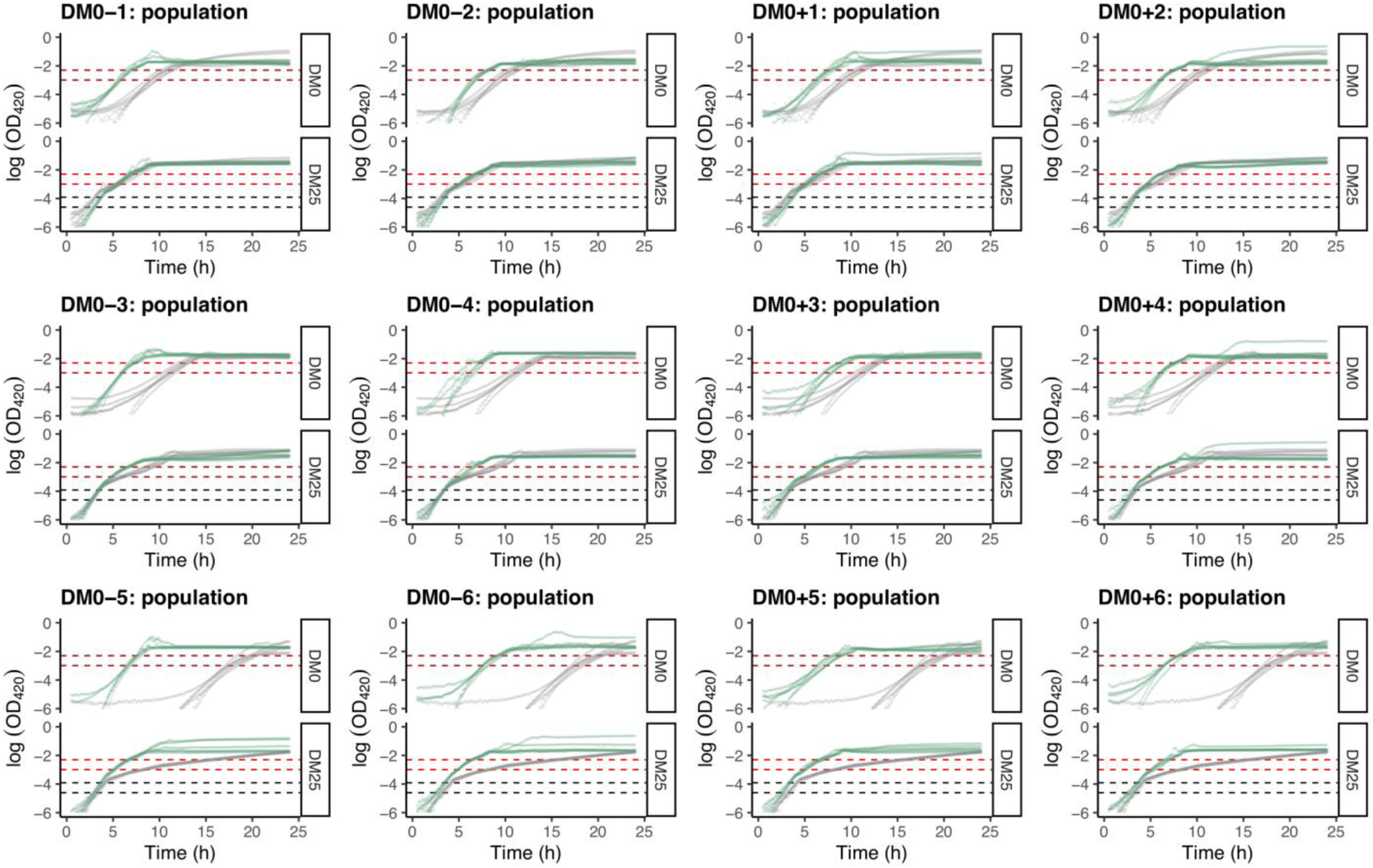
Log*_e_*-transformed growth curves of the 12 DM0-evolved whole-population samples, measured in DM0 and DM25. Dashed black and red lines indicate the intervals used to calculate growth rates on glucose and citrate, respectively (Supplementary Fig. S1). See Supplementary Figure S3 for additional details.

**Figure S5.**
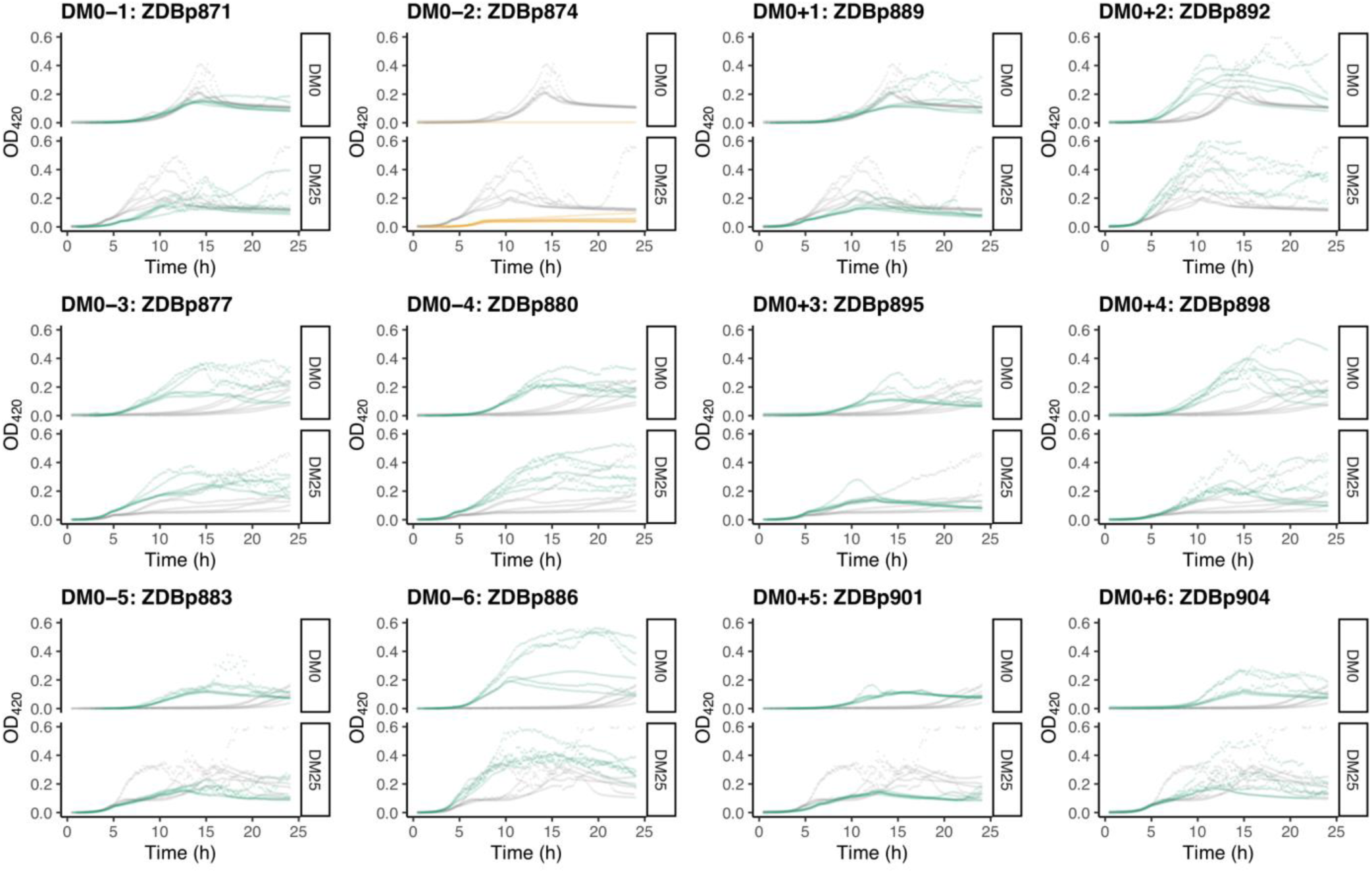
Growth curves of the 12 DM0-evolved clones, measured in DM0 and DM25. For comparison, growth curves of the evolved clones are paired with those of their ancestors: CZB151 (top row), CZB152 (middle row), and CZB154 (bottom row). The evolved and ancestral curves are shown in green and gray, respectively, except the anomalous Cit^−^ evolved clone is shown in orange.

**Figure S6.**
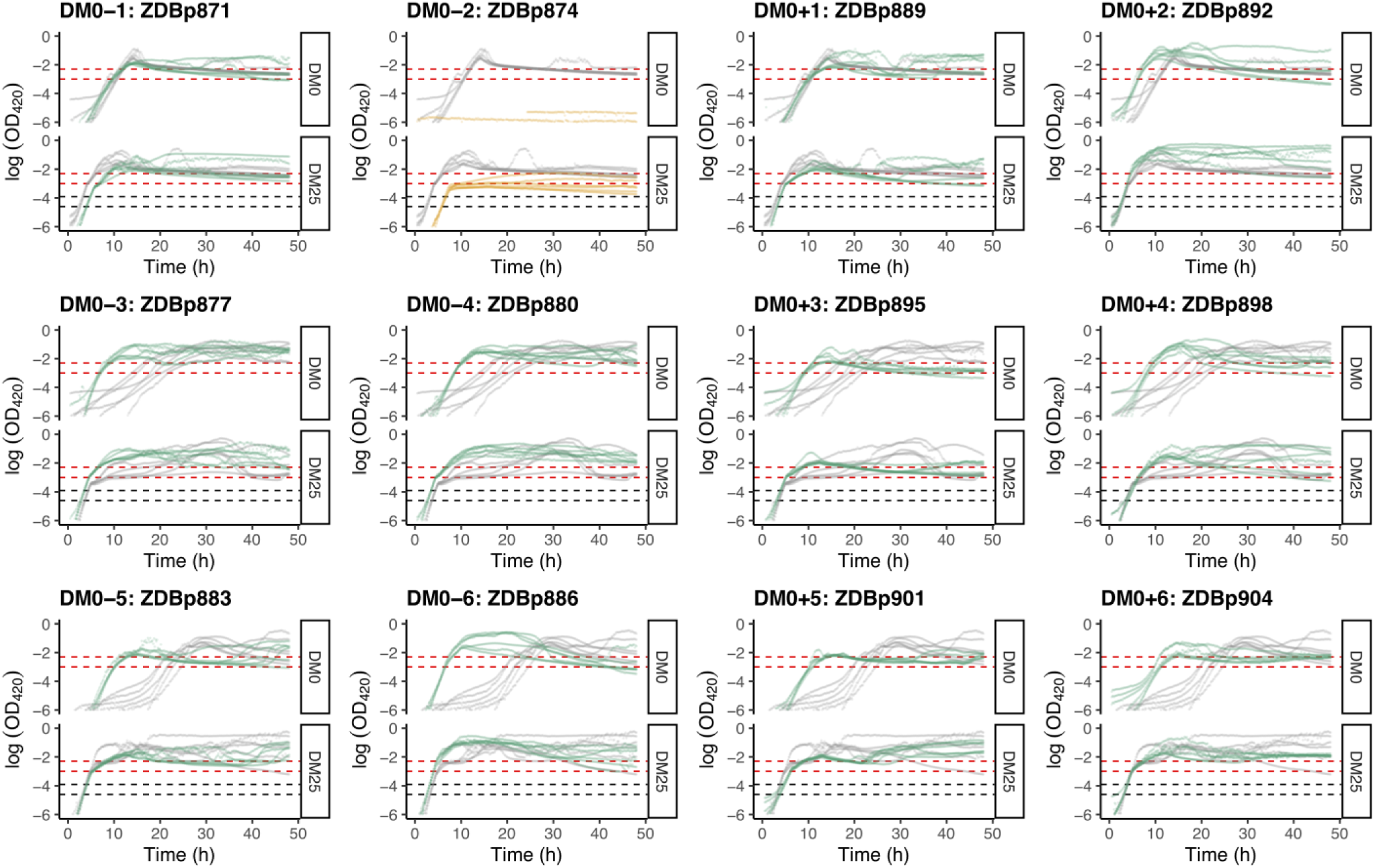
Log*_e_*-transformed growth curves of the 12 DM0-evolved clones, measured in DM0 and DM25. Dashed black and red lines indicate the intervals used to calculate growth rates on glucose and citrate, respectively (Supplementary Fig. S1). See Supplementary Figure S5 for additional details.

**Figure S7.**
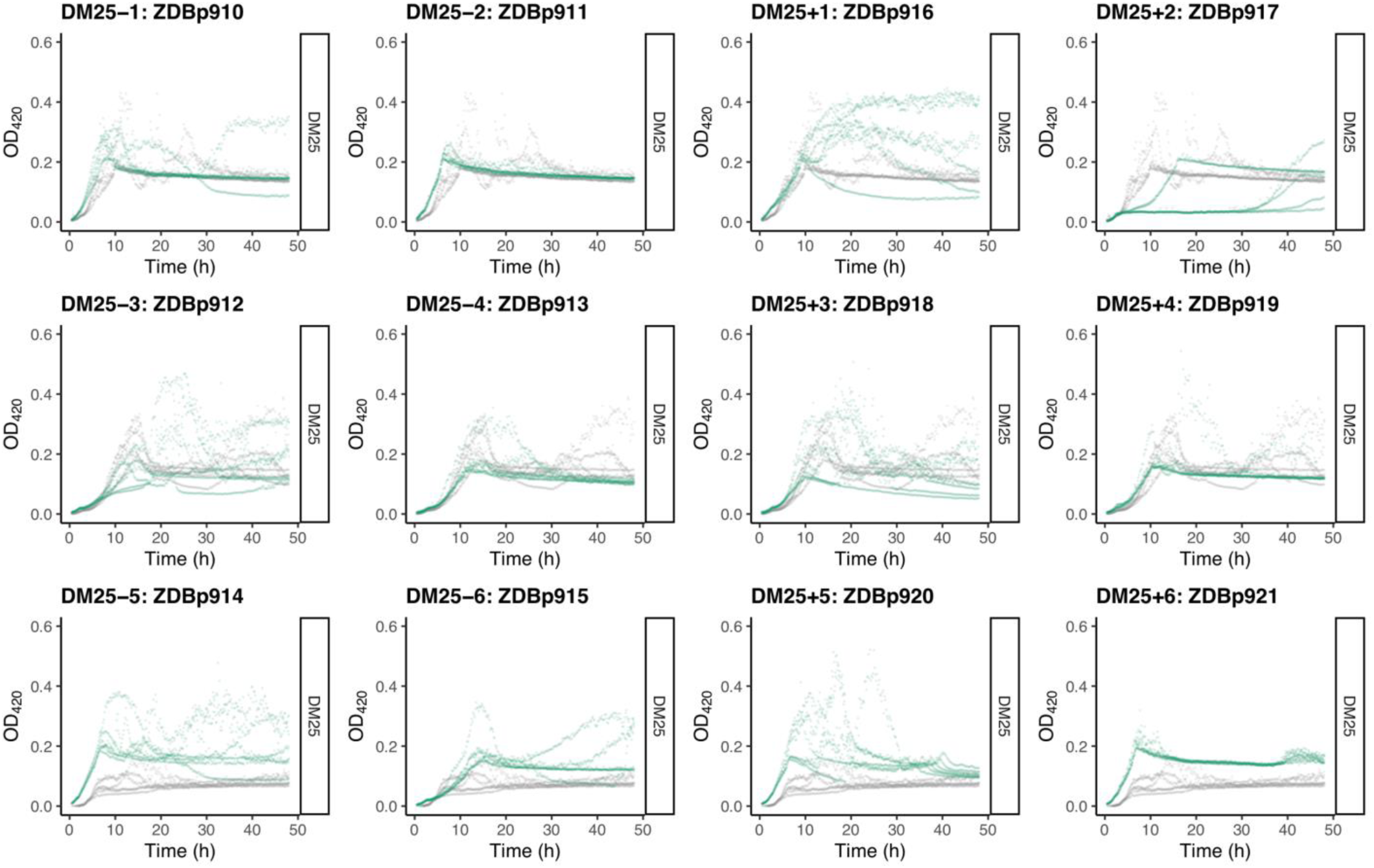
Growth curves of the 12 DM25-evolved clones, measured in DM25 only. (Many DM25-evolved clones grew inconsistently in DM0.) For comparison, growth curves of the evolved clones are paired with those of their founders: CZB151 (top row), CZB152 (middle row), and CZB154 (bottom row). The evolved and ancestral curves are shown in green and gray, respectively.

**Figure S8.**
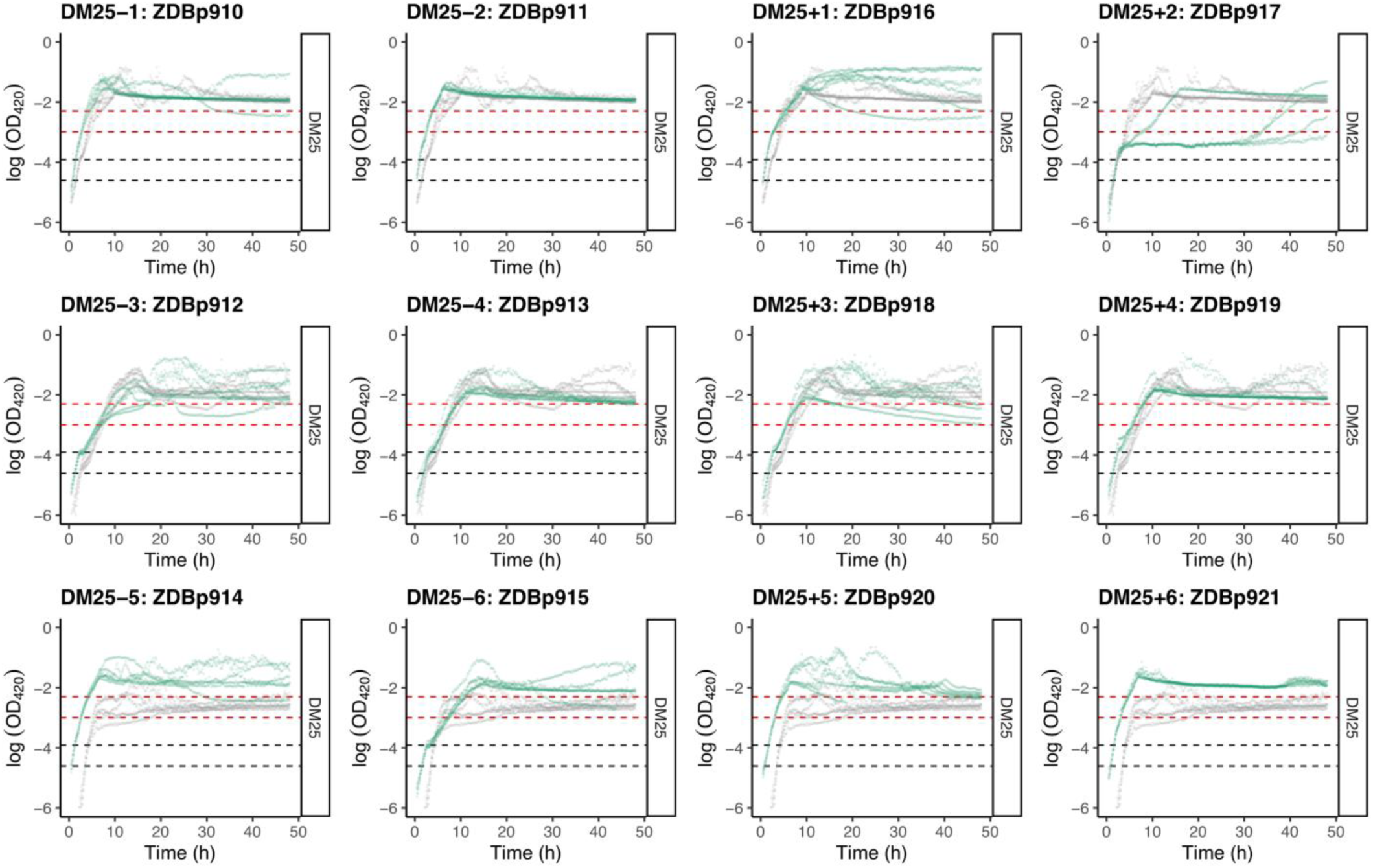
Log*_e_*-transformed growth curves of the 12 DM25-evolved clones, measured in DM25. Dashed black and red lines indicate the intervals used to calculate growth rates on glucose and citrate, respectively (Supplementary Fig. S1). See Supplementary Figure S7 for additional details.

**Figure S9.**
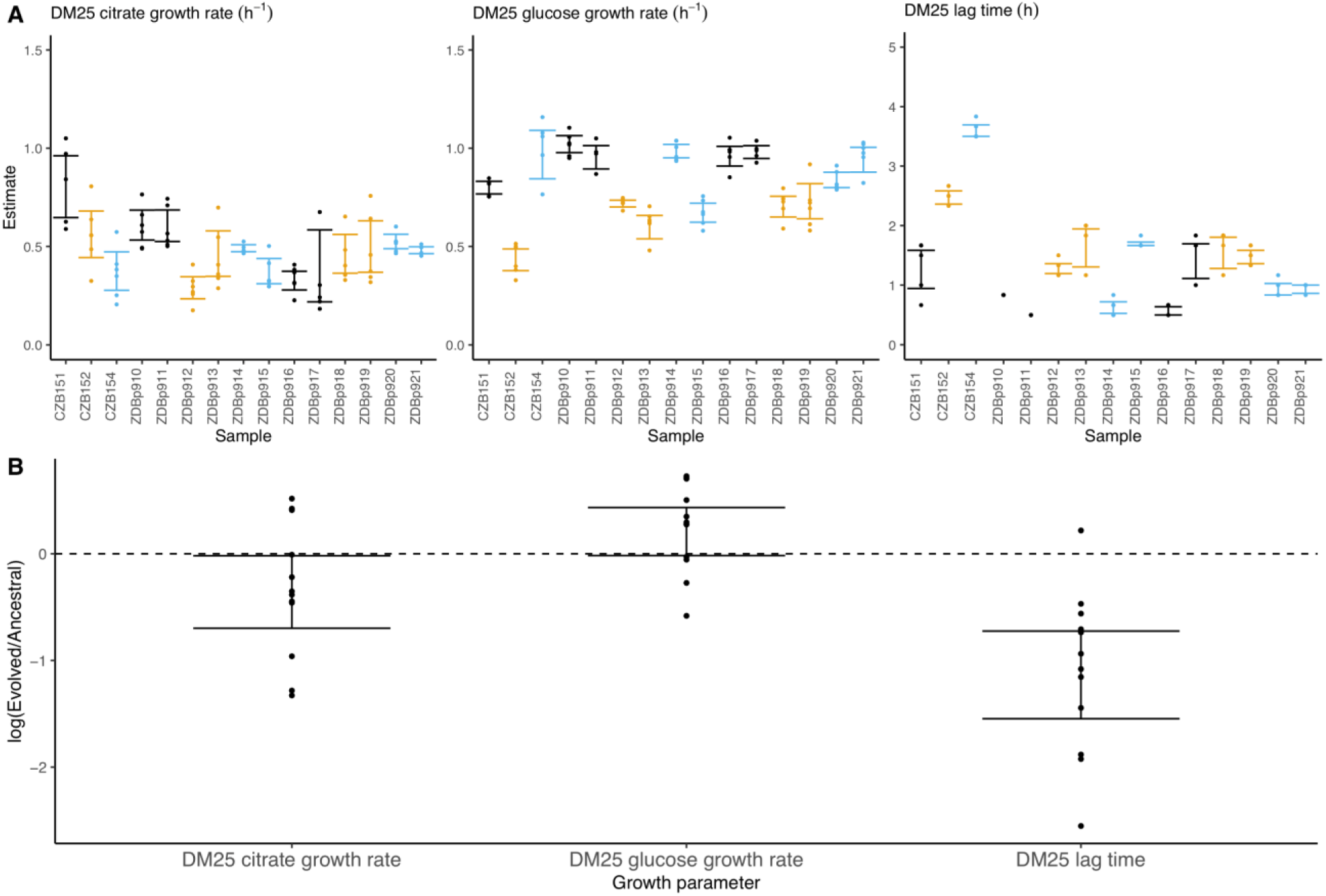
Growth parameters of the 12 DM25-evolved clones and their 3 Cit^+^ ancestors. A) Estimates of growth parameters for each ancestral and DM25-evolved clone, using the log-slope method (Supplementary Fig. S1). Estimates for ancestral strain CZB151 and its descendants are shown in black, estimates for CZB152 and its descendants are in orange, and estimates for CZB154 and its descendants are in blue. Units for growth rates *r* are h^−1^, and units for lag times are h. Bias-corrected and accelerated (*BC_a_*) bootstrap 95% confidence intervals around parameter estimates were calculated using 10,000 bootstraps; no confidence interval is shown if a parameter could not be estimated accurately from the available data. Aberrant estimates that fall outside of these ranges are not shown. B) Estimates of log_2_-transformed ratios of growth parameters for the evolved clones and their ancestors. The growth curves used to estimate these parameters are shown in Supplementary Figures S7 and S8.

**Figure S10.**
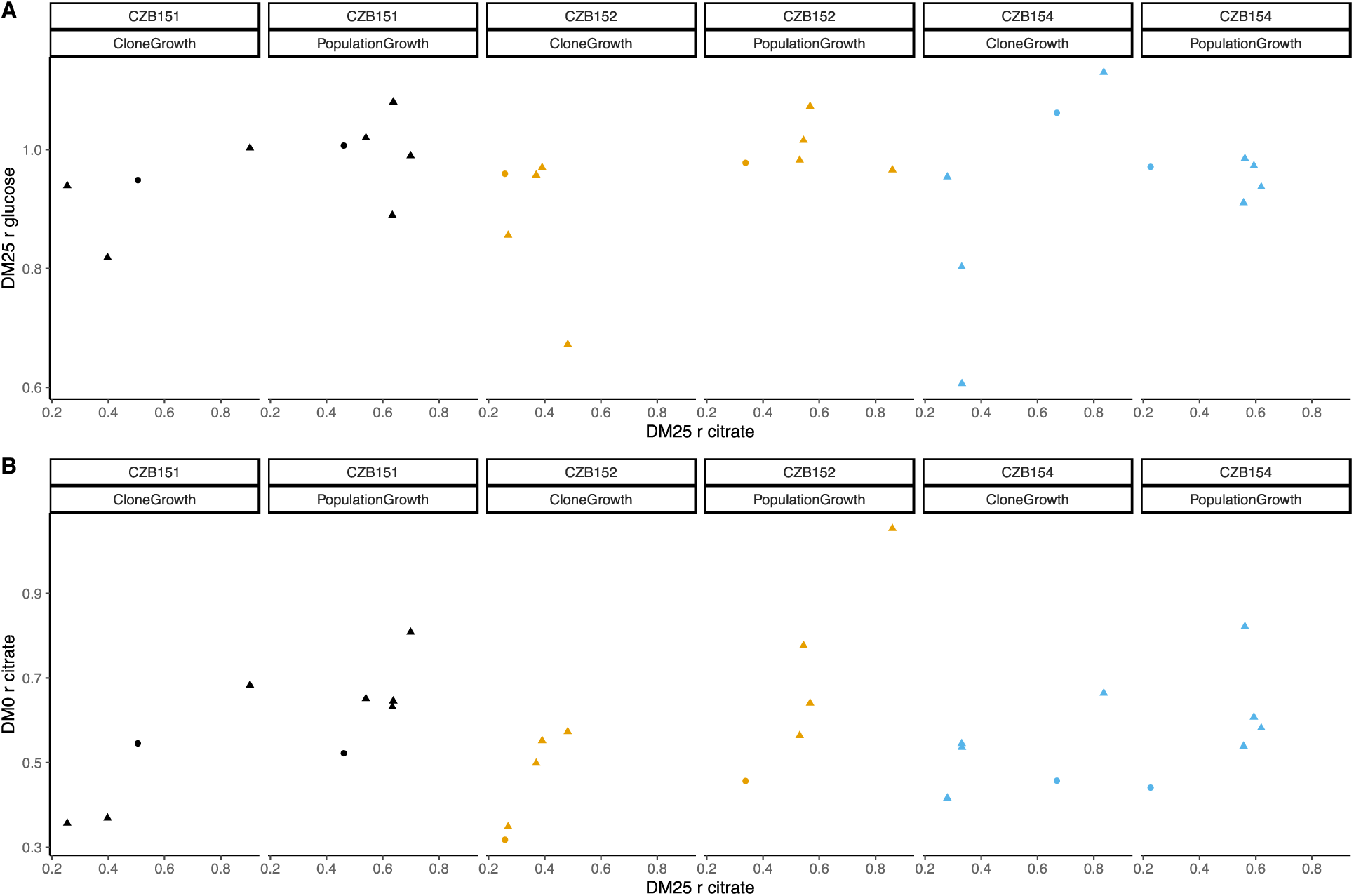
Correlations between estimated growth rates across substrates and media. All tests are two-tailed, because growth rates across substrates and media might, in principle, exhibit tradeoffs. A) Correlations between *r_glucose_* and *r_citrate_* in DM25 are not significant (Pearson’s *r* = 0.4788, d.f. = 12, *p* = 0.0833 for clones; *r* = –0.0392, d.f. = 13, *p* = 0.8897 for populations). B) Correlations between *r_citrate_* in DM0 and *r_citrate_* in DM25 are highly significant (*r* = 0.7513, d.f. = 12, *p* = 0.0020 for clones; *r* = 0.8041, d.f. = 13, *p* = 0.0003 for populations). Circles and triangles indicate ancestral and evolved samples, respectively. Colors distinguish the different Cit^+^ ancestors and their evolved descendants.

**Figure S11.**
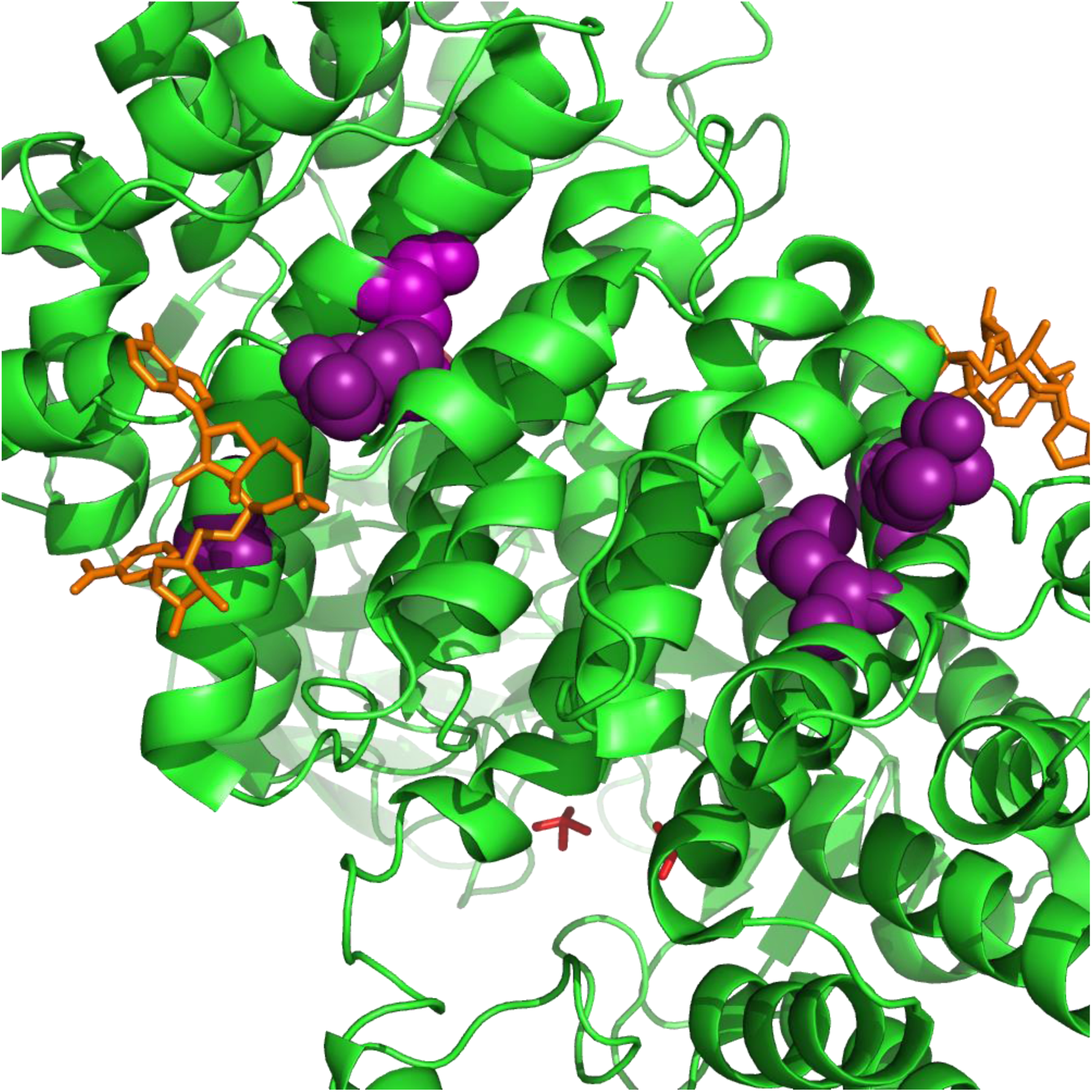
Parallel substitutions at the amino-acid level in citrate synthase, GltA. All of the evolved substitutions occur at the allosteric protein-ligand interface with NADH. GltA is shown in its dimeric, NADH-bound conformation (1NXG crystal structure in the Protein DataBank). The M172I, A162T, I114F substitutions are shown in purple. NADH is shown in orange.

**Figure S12.**
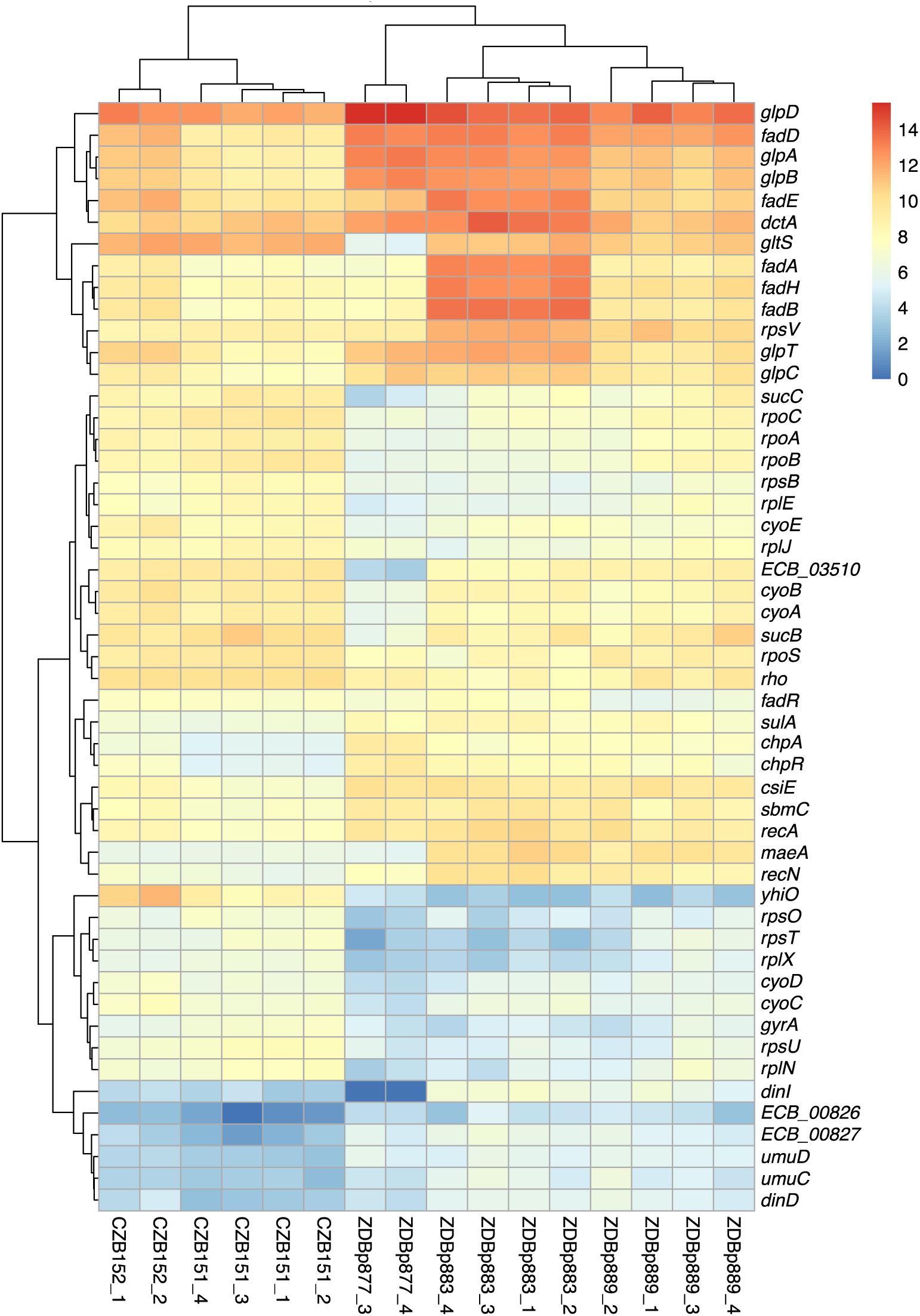
Transcriptomic analysis of ancestral and evolved clones. Differential expression analysis comparing two ancestral (CZB151 and CZB152) and three evolved clones (ZDBp877, ZDBp883, ZDBp889), produced by *sleuth* (Pimentel *et al*. 2017). The colored bar (at right) shows the level of RNA expression based on estimated counts and transformed as log_2_(1 + est_counts). The differentially expressed genes discussed in the main text are shown here. The numerical labels after the strain identifiers indicate the 2 or 4 biological replicates for each clone (i.e., RNA samples prepared from independently revived cultures of that clone).

